# Targeting the CaVα-β interaction yields a selective antagonist of the N-type CaV2.2 channel with broad antinociceptive efficacy

**DOI:** 10.1101/492181

**Authors:** Aude Chefdeville, Jie Yu, Xiaofang Yang, Aubin Moutal, Vijay Gokhale, Zunaira Shuja, Lindsey A. Chew, Shreya S. Bellampalli, Shizhen Luo, Liberty François-Moutal, Taehwan Ha, Samantha Perez-Miller, Ki Duk Park, Amol M. Patwardhan, John M. Streicher, Henry M. Colecraft, May Khanna, Rajesh Khanna

**Author notes:** co-first authors (Aude Chefdeville, Jie Yu). Corresponding Author*: Dr. Rajesh Khanna, Department of Pharmacology, College of Medicine, University of Arizona, 1501 North Campbell Drive, P.O. Box 245050, Tucson, AZ 85724, USA.

## Abstract

Inhibition of voltage-gated calcium (CaV) channels is a potential therapy for many neurological diseases including chronic pain. Neuronal CaV1/CaV2 channels are composed of α, β and α2δ subunits. The β-subunits of CaV channels are cytoplasmic proteins that increase the surface expression of the pore-forming α subunit of CaV. We targeted the high-affinity protein-protein interface of CaVβ’s pocket within the CaVα-subunit. Structure-based virtual screening of 50,000 small molecule library docked to the β-subunit led to the identification of 2-(3,5-dimethylisoxazol-4-yl)-N-((4-((3-phenylpropyl)amino)quinazolin-2-yl)methyl)acetamide (compound ***45***). This small molecule bound to CaVβ and inhibited its coupling with N-type voltage-gated calcium (CaV2.2) channels, leading to a reduction in CaV2.2 currents in rat dorsal root ganglion (DRG) sensory neurons, decreased pre-synaptic localization of CaV2.2 *in vivo*, decreased frequency of spontaneous excitatory post-synaptic potentials (sEPSC), and inhibited release of the nociceptive neurotransmitter calcitonin gene related peptide (CGRP) from spinal cord. ***45*** was antinociceptive in naïve animals and reversed allodynia and hyperalgesia in models of acute (post-surgical) and neuropathic (spinal nerve ligation, chemotherapy- and gp120-induced peripheral neuropathy, and genome-edited neuropathy) pain. ***45*** did not cause akinesia or motor impairment, a common adverse effect of CaV2.2 targeting drugs, when injected into the brain. ***45***, a quinazoline analog, represents a novel class of CaV2.2-targeting compounds that may serve as probes to interrogate CaVα-β function and ultimately be developed as a non-opioid therapeutic for chronic pain.

## 1. Introduction

The N-type voltage-gated calcium (CaV2.2) channels are critical determinants of increased neuronal excitability and neurotransmission accompanying persistent neuropathic pain [6; 9; 12; 65]. Expressed in the presynaptic termini of primary afferent nociceptors in the spinal cord [57], CaV2.2 represent a control point for synaptic activity. The potential of targeting CaV2.2 has been demonstrated with attenuation of allodynia by CaV2.2-blocking conotoxins [49] and the altered pain behavior of CaV2.2 knock-out mice [26; 27]. CaV2.2 remains a high-value target with several companies developing CaV2.2-targeted compounds [29; 64]. Ziconotide (Prialt®)[33] and Gabapentin (Neurontin®) directly target different elements of the CaV2.2 complex. These drugs, however, present with problematic side effects, difficult dosing regimens and have high number needed to treat (NNT) values [16]. Therefore, the development of novel CaV2.2-targeted drugs with improved efficacy and therapeutic index is highly desirable [43]. We have advanced a strategy targeting protein interactions that modulate CaV2.2 as an alternative to direct channel blockade [7; 15; 17; 59; 60]. Targeting *channel regulation* may potentially lessen many of the adverse side effects associated with *direct* channel block. Supporting this concept, we reported that targeting CaV2.2 *indirectly* with a peptide derived from an ancillary regulatory protein did not affect memory retrieval, motor coordination, depression-associated behaviors [7] and was not rewarding/addictive [17].

Here, we focused on the β subunits of CaV channels. CaVβs are cytoplasmic proteins encoded by four different genes (β_1-4_), including multiple splice variants [14]. Their signature roles are to increase the surface expression of the pore-forming α subunit of Ca^2+^ channels [14; 45] and regulate biophysical properties of the α subunit of Ca^2+^ channels [14; 45]. The sum effect of these ancillary roles is to increase the amount of Ca^2+^ influx within cells. In neurons, Ca^2+^ influx triggers neurotransmitter release, where the amount of transmitter released from a presynaptic terminal is dependent on the amount of Ca^2+^ entering the terminal [3; 47]. Further involvement of β subunits in regulating synaptic transmission is also supported by evidence of protein-protein interactions of the CaVβ subunits with the synaptic vesicle release machinery [54]. Thus, the subunit is part of the transmitter release site core complex, at the center of which resides the α subunit of the Ca^2+^ channel [25]. Altered CaVβ subunit expression will modulate the function of the α subunit of the calcium channel and can underlie pathological neuronal transmission. In neuropathic pain, CaVβ3 expression is increased and leads to augmented high-voltage-gated Ca^2+^ channel function in small diameter sensory neurons [30]. This pathological increase of CaVβ3 amplifies spinal nociceptive neurotransmission and is sufficient to induce pain [30]. Thus, pharmacotherapeutic approaches targeting the CaVβ/CaVα interface could restore physiological Ca^2+^ homeostasis and be beneficial for chronic pain management.

We utilized a computational approach to target a protein-protein interface important for voltage-gated calcium channel activity. We report the identification of the small molecule 2-(3,5-dimethylisoxazol-4-yl)-N-((4-((3-phenylpropyl)amino)quinazolin-2-yl)methyl)acetamide (***45***) targeting the CaVβ/CaVα interface. Here, we demonstrate that ***45*** (i) specifically inhibits CaV2.2 (over other subtypes) in sensory neurons, (ii) acts at presynaptic sites, (iii) blunts release of the pro-nociceptive neurotransmitter CGRP, (iv) reverses thermal hyperalgesia and mechanical allodynia in acute and neuropathic models of pain, and (v) has no effect on motor activity. This quinazoline analog may be used to mitigate chronic pain by controlling CaV2.2 function.

## 2. Materials and Methods

Detailed descriptions of methods used, and any associated references are available in SI Materials and Methods. Briefly, all biochemical, electrophysiology and behavior experiments were performed in a blinded fashion according to established protocols [7; 37; 40]. All animal protocols were approved by the Institutional Animal Care and Use Committee of the College of Medicine at the University of Arizona and conducted in accordance with the Guide for Care and Use of Laboratory Animals published by the National Institutes of Health.

## 3. Results

### 3.1. A quinazoline compound (45) targets the CaVα-CaVβ interface and inhibits depolarization evoked Ca^2+^ influx in sensory neuron subpopulations

To identify small molecules that disrupt the CaVα–β interaction, we used a virtual screening approach against the crystal structure of the complex formed between the CaVβ2a subunit and a peptide of the α1c subunit (*Fig. 1A*) [52]. The alpha interaction domain (AID) peptide was removed and its binding site was used for docking 50K drug-like small molecules (molecular weight ≤ 500 Da) commercially available from ChemBridge Inc. The resulting complexes were ranked using Glide score and other energy-related terms [18]. Forty-nine compounds docked into the α-binding pocket on CaVβ2a. The 49 compounds were screened by Ca^2+^ imaging for their ability to inhibit depolarization-induced calcium influx in rat DRG neurons. Of these compounds, 13 were either insoluble or killed the neurons, and 11 compounds inhibited Ca^2+^ influx by ≥50%. One docked compound, ***45***, engaged all three (V241, I343, N390) AID “hotspot” residues (*Fig. 1B*). The structure of ***45*** aligned with 3 residues (W440, I441, Y437) from the AID (*Fig. 1C, E*) and specifically bound CaVβ2a protein in saturation transfer difference nuclear magnetic resonance (STD-NMR) (*Fig. 1D*).

**Figure 1.**
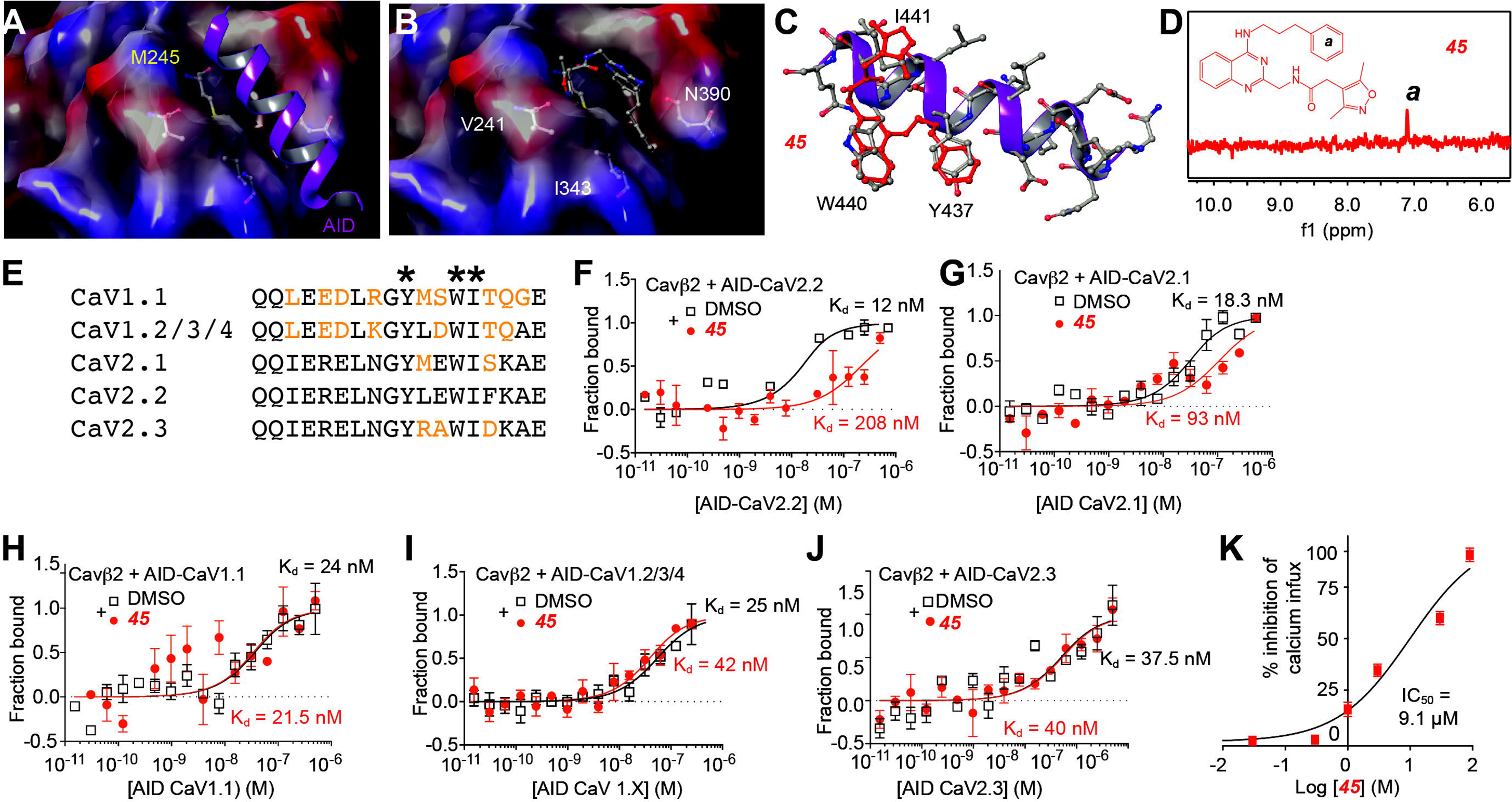
Structure-guided high-throughput screen of CaV α-β interaction inhibitors identifies 45. (*A*) Space-filling model of the CaVβ2a subunit showing its α-binding pocket (ABP). A ribbon representation of the CaV α-interaction domain (AID, purple) and the AID’s key interaction residue (M245) are highlighted. (*B*) Ball-and-stick representation of key ABP residues are shown (i.e., V241, I343, and N390). (*C*) A three-dimensional representation of ***45*** (red) overlaid on the AID’s reveals shared chemical features, especially at AID residues W440, I441, and Y437. (*D*) 1D ^1^H STD NMR showing on-resonance difference spectrum for ***45*** (red); only regions that yielded a STD signal in the presence of ***45*** are plotted. The two-dimensional chemical structure of compound ***45*** shows that region *a* corresponds with the STD signal highlighted. (*E*) Amino acid alignments of AID residues of the indicated calcium channel subtypes. Residues different from the CaV2.2 core AID sequence are denoted in orange. Asterisks denote the key residues conserved in all AID sequences involved in the interaction with the beta subunits. Microscale thermophoresis (MST) was used to determine dissociation constants for binding of AID-CaV2.2 (*F*), CaV2.1 (*G*), CaV1.1 (*H*), CaV1.2/3/4 (*I*) (500-0.01nM), and CaV2.3 (*J*) to Beta2-CaVβ2-His (25nM) in the presence (filled red circles) or absence of 10 μM ***45*** (open squares). Data is presented as means ± SEM. (*K*) Concentration-response curve illustrates that ***45*** inhibits membrane depolarization-evoked Ca^2+^ influx in DRGs (n>150 cells per concentration from 3 independent platings) in a concentration dependent manner.

Microscale thermophoresis – a method that permits analysis of biomolecular interactions, demonstrated that ***45*** reduced the binding of CaVβ2-His to the AID-CaV2.2 peptide from a K_d_ of 12 nM to 208 nM (Fig. 1F) and reduced the binding of CaVβ2-His to the AID-CaV2.1 peptide from a K_d_ of 18.3 nM to 93 nM (*Fig. 1G*) while binding of CaVβ2-His to the AID peptides from CaV1.1 (*Fig. 1H*), CaV1.2/CaV1.3/CaV1.4 (*Fig. 1I*), and CaV2.3 (*Fig. 1J*) were unaffected. Whether the inhibition was achieved through 45 binding to the predicted pocket on CaVβ or through allosteric modulation remains to be determined.

***45*** inhibited depolarization-evoked Ca^2+^ influx in a concentration-dependent manner in DRG sensory neurons with an IC50 of ~9.1μM (*Fig. 1K*). Since compounds are known to aggregate at screening-relevant concentrations in every compound library, we queried the Aggregator Advisor database [13] with ***45***, which revealed no similarity to known aggregators. A similar query of ***45*** in the Zinc15 database revealed no hits to molecules containing PAINS chemotypes. As the Aggregator Advisor calculated a partition coefficient for ***45*** in n-octanol/water (cLogP) value of 3.5, which is in the range reported for many other aggregators [22], dynamic light scattering was performed. Dynamic light scattering curve of ***45*** in the presence of a non-ionic detergent (0.1% Tween-20) did not reveal significant colloidal aggregation with only less than 9% of the compound forming particles around a 50 nm radius (*Fig. S1A*). To further control for potential effects of aggregation, we subjected ***45*** to a centrifugation spin-down (15 min at 21,000 X g) and then used the supernatant from the spin-down of ***45*** to perform calcium imaging studies in sensory neurons. There were no differences in the depolarization-triggered calcium responses between cells treated with ***45*** irrespective of whether it was centrifuged or not (*Fig. S1B*). These data, along with the use of non-ionic detergent in the MST experiments (*Fig. 1E, F*) as well as Tween-80 in the vehicle for all *in vivo* experiments triangulate to eliminate colloidal aggregation as a mechanism of ***45***’s inhibitory effect. Ca^2+^ influx was inhibited by ***45*** in all classes of DRG neurons (*Fig. S2A-F*), identified by constellation pharmacology – a method that permits functional ‘fingerprinting’ of neurons to specific receptor agonists [51].

### 3.2. Lack of activity of 45 for opioid receptors

Inhibition of Ca^2+^ influx in sensory neurons can occur via activation of opioid receptors [23]. Consequently, we tested whether opioid receptors could be engaged by ***45*** by competition radioligand binding at all 3 opioid receptors *in vitro*. We competed ***45*** and a positive control compound (naloxone for MOR and DOR, U50,488 for KOR) versus a fixed concentration of ^3^H-diprenorphine in Chinese Hamster Ovary (CHO) cells expressing the human μ (MOR), δ (DOR), or κ (KOR) opioid receptor. ***45*** (up to 10 μM) did not bind to the MOR or KOR, and only bound to the DOR with very weak affinity (*Fig. S2G-I*). In contrast, the positive control compounds bound to all 3 targets with expected affinity (*Fig. S2G-I*). Thus, inhibition of Ca^2+^ influx by ***45*** likely does not involve opioid receptors. Compound ***45*** was evaluated at UNC’s NIMH Psychoactive Drug Screening Program (PDSP) against a battery of 30 receptors known to adversely impact drug effectiveness. No significant binding was observed at 20 μM. Moreover, ***45*** did not affect hERG K^+^ channel activity at 20 μM, which helps to confirm drug safety (data not shown).

### 3.3. Preferential activity of 45 for CaV2.2

Because the CaVα–β pocket is conserved across the various α and β subunits, we next set out to determine if ***45*** was selective for any of the isoforms. Using whole-cell patch-clamp electrophysiology, we measured Ca^2+^ currents in DRG neurons treated with 20 µM of ***45*** (~twice the IC_50_ of ~9.1μM) or 0.1% DMSO as a control (*Fig. 2A*). Consistent with our Ca^2+^ screening, ***45*** inhibited (by ~50%) total Ca^2+^ currents in DRG sensory neurons (*Fig. 2B, C*), without affecting the biophysical properties (activation and inactivation) of these channels (*Fig. 2D, E*). Next, using saturating concentrations of selective Ca^2+^ channel inhibitors, we isolated either CaV1 (L-type), CaV2.1 (P/Q-type), CaV2.2 (N-type) or CaV2.3 (R-type) Ca^2+^ currents in DRGs to evaluate the inhibitory potential of ***45*** on each subtype. ***45*** (20 μM) inhibited only N-type Ca^2+^ channels (>70%) but had no effect on the other subtypes (*Fig. 2F*). Next, we expressed the α-subunit of CaV2.2 in combination with CaVβ_1-4_ subunits in heterologous cells and measured whole cell currents. ***45*** inhibited CaV2.2 currents when co-expressed with CaVβ subunits 1-3 (*Fig. S3A-C*), but not with CaVβ4 (*Fig. S3D*). Together, these results demonstrate selectivity of ***45*** for N-type channels with CaVβ subunits 1-3, but not CaVβ4.

**Figure 2.**
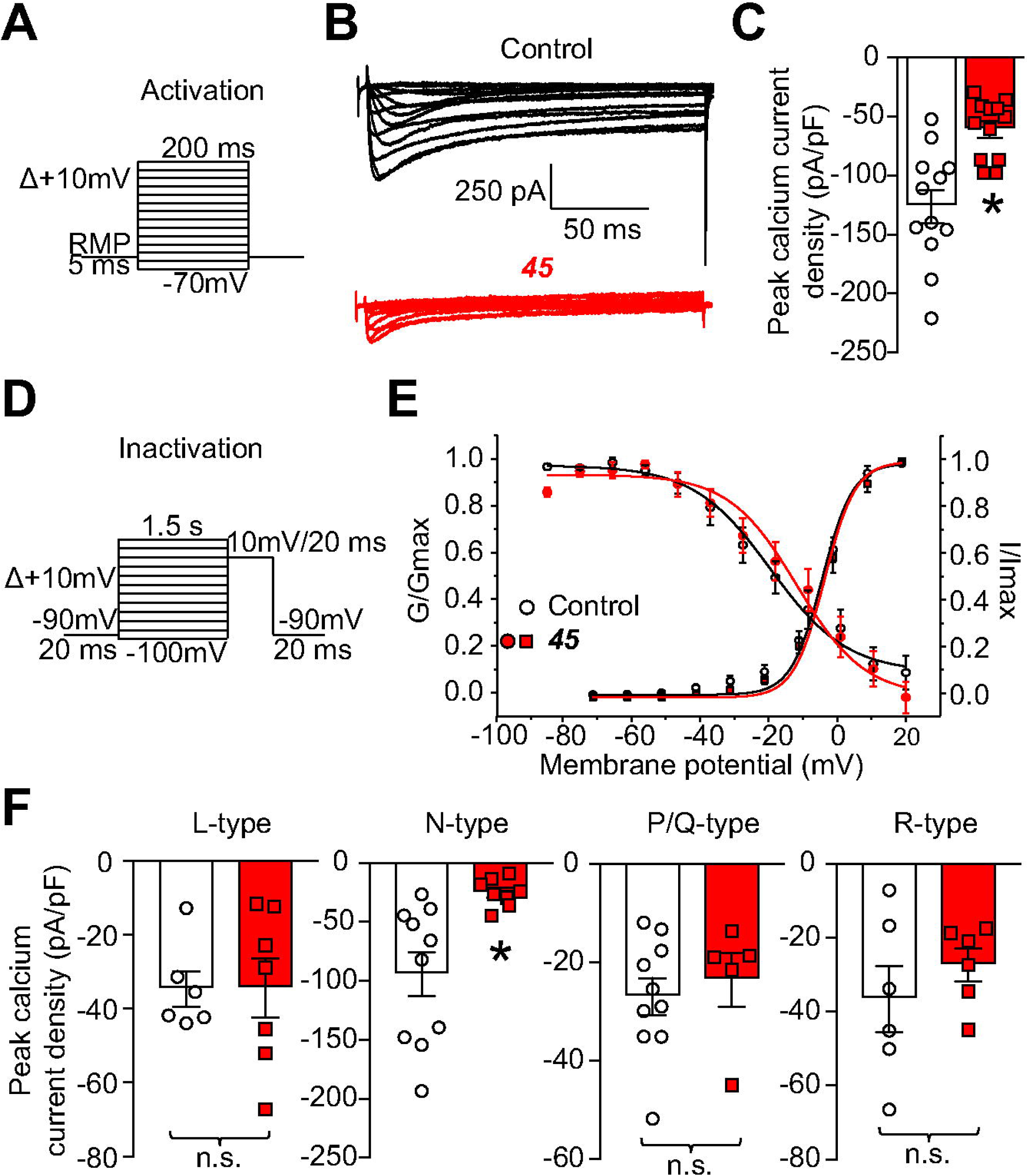
*45* selectively inhibits Ca^2+^ currents mediated by CaV2.2. (*A*) Activation (i.e., current-voltage (I-V) relationship voltage step protocol). Cells were held at resting membrane potential for 5-ms before depolarization by 200-ms voltage steps from −70 mV to +60 mV in 10 mV increments. Currents were normalized to each cell’s capacitance (pF). This allowed for collection of current density data to analyze activation of Ca^2+^ channels as a function of current vs. voltage as well as peak current density. (*B*) Representative Ca^2+^ current traces from DRGs subjected to the activation protocol (shown in A). (*C*) Summary graph of peak Ca^2+^ current density (pA/pF) from DRGs incubated with 0.1% DMSO or 30µM ***45*** overnight (n=12; *P < 0.05, Kruskal–Wallis test with Dunnett’s post hoc comparisons). (*D*) Inactivation voltage step protocol. Cells were held at −90 mV for 20-ms before depolarization by 1.5-s voltage steps from −100 mV to +10 mV in 10 mV increments, followed by a 20-ms pulse at 10 mV before returning to −90 mV for 20-ms. (*E*) Normalized peak current plotted against its preceding holding potential and fitted with the Boltzmann relation. No significant differences were detected in half-maximal voltage or slope properties of either activation or inactivation between DMSO and ***45*** conditions. (*F*) Summary graph of peak Ca^2+^ current density (pA/pF). Presence of ***45*** did not significantly reduce Ca^2+^ currents pharmacologically isolated through L-type, P/Q-type, or R-type Ca^2+^ channels. Presence of ***45*** significantly reduced N-type Ca^2+^ currents (compared to DMSO treated DRGs, n=6-10; *P < 0.05, Kruskal–Wallis test with Dunnett’s post hoc comparisons).

### 3.3. 45 inhibits spinal neurotransmission

CaV2.2 is involved in spinal nociceptive neurotransmission [20] through the control of pre-synaptic nociceptive neurotransmitter release from C-fibers [7; 20; 38; 50]. ***45***-mediated inhibition of CaV2.2 from DRG neurons should in turn decrease spontaneous excitatory post-synaptic currents (sEPSCs) measured in the *substantia gelatinosa* of the dorsal horn of the spinal cord. We recorded sEPSCs from neurons in laminae I-II in this region of the lumbar dorsal horn (*Fig. 3A, B*). No change was observed in the series resistance of cells perfused with ***45*** (20 µM) compared with control (*Fig. 3C*). ***45*** treatment had no effect on the amplitude of the recorded sEPSCs (*Fig. 3D-E*). We observed a decrease of sEPSC frequency after treatment with 20 µM of compound ***45*** (*Fig. 3F-G*). Next, we recorded EPSCs evoked by a paired pulse stimulation protocol, which is commonly used to assess changes in presynaptic function [66; 67]. At 50-ms intervals, paired-pulse ratios were increased by ***45*** treatment (*Fig. 3H-I*). These results collectively implicate a presynaptic provenance for the mechanism of ***45***’s action.

**Figure 3.**
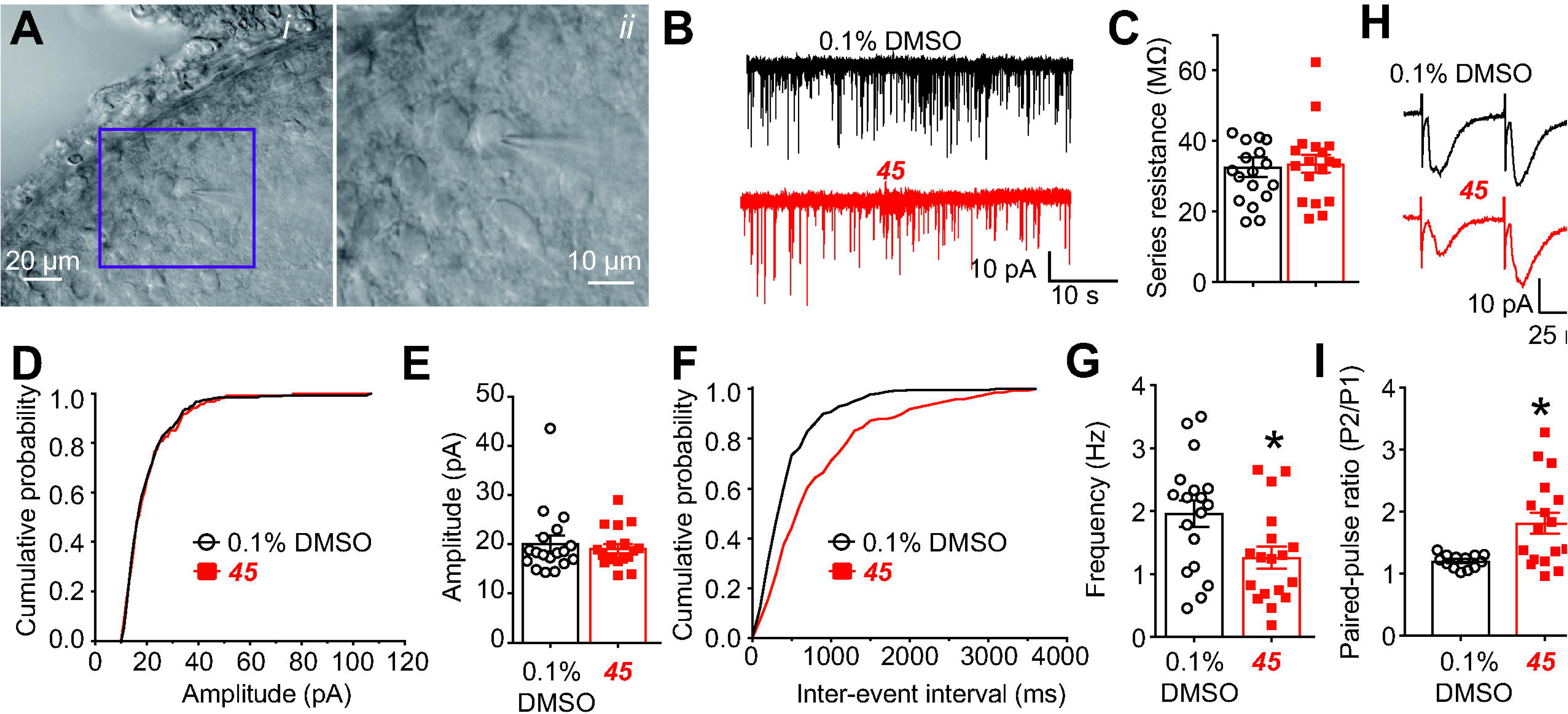
*45* decreases spontaneous excitatory synaptic transmission. Photomicrograph of slice preparation (showing that the substantia gelatinosa (SG) can be identified as a translucent pale band in the superficial dorsal horn enabling positioning of the recording electrode to this region. (A, *i*) infrared differential interference contrast image, and (A, *ii*) image of the same cell (indicated by a purple red box in middle panel) with part of the recording electrode after whole-cell configuration. (B) Representative recording traces of cells from the indicated groups: 0.1% DMSO or 20 μM ***45***. (C) Series resistance in unchanged between the two conditions. The cumulative probability of amplitude (D) and inter-event interval (F). Summary of amplitudes (E) and frequencies (G) of sEPSCs for the indicated groups are shown. Data are expressed as means ± SEM from n= 17-18 cells per condition. *p < 0.05 (versus DMSO); Student’s t-test. (H) Representative traces of paired pulse ratio (PPR) from the indicated groups: 0.1% DMSO or 20 μM ***45***. (I) Summary of average PPRs from the indicated groups. Data are expressed as means ± SEM from n= 13-17 cells per condition. *p < 0.05 (versus DMSO); Student’s t-test.

### 3.4. 45 reduces CaV2.2 spinal presynaptic localization

***45*** can inhibit CaV2.2 AID interaction with the beta subunits. We asked if interfering with this interaction using ***45*** in vivo could affect CaV2.2 presynaptic localization in lumbar dorsal horn of the spinal cord. We first extracted synaptosomes from the dorsal horn of the spinal cord of rats 1 hour after injection with ***45*** (2µg in 5µl, i.th.) and isolated pre- and post-synaptic fractions (*Fig. 4A*). The fractionation efficiency was verified by western blot where the post-synaptic marker PSD95 was highly enriched in the PSD fraction while the pre-synaptic marker synaptophysin was only found in the non-PSD fraction (*Fig. 4A*). Indeed, CaV2.2 was exclusively localized in the pre-synaptic fraction (*Fig. 4A*). We then focused on the pre-synaptic (non-PSD) fraction to evaluate if ***45*** could change the pre-synaptic levels of CaV2.2. We found that ***45*** decreased CaV2.2 pre-synaptic levels (*Fig. 4B, C*). Notably, ***45*** did not affect the localization of the NaV1.7 voltage-gated sodium channel NaV1.7 (*Fig. 4B, C*). These results demonstrate that interfering with the CaV2.2 AID/beta-subunit interaction specifically reduces the pre-synaptic localization of the CaV2.2 *in vivo*.

**Figure 4.**
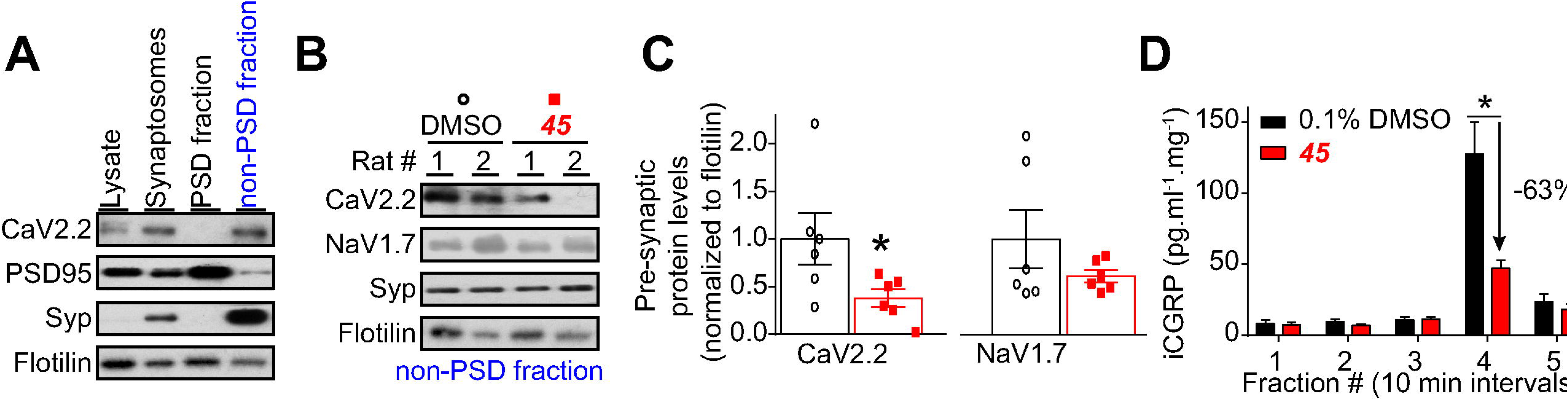
*45* decreases CaV2.2 pre-synaptic localization in vivo and reduces evoked calcitonin gene related peptide (CGRP) release from spinal cord. (*A*) Immunoblots showing the integrity of the synaptic fractionation from lumbar dorsal horn of the spinal cord. The non-post synaptic density (non-PSD) fraction was enriched in the pre-synaptic marker Synaptophysin (Syp) and the PSD fraction was enriched in the post-synaptic marker PSD95. The membrane-associated protein flotillin was used as a loading control. (*B*) Immunoblots showing the presynaptic CaV2.2 levels in the lumbar dorsal horn of the spinal cord of animals having received ***45*** (2µg in 5 µl, i.th.) compared to vehicle (0.1% DMSO). Synaptophysin shows the integrity of each fraction. Flotilin is used as a loading control. (*C*) Bar graph showing decreased CaV2.2 levels at the presynaptic sites of lumbar dorsal horn of the spinal cord in ***45*** treated animals. Mean ± S.E.M., *p<0.05, Mann-Whitney compared to the DMSO vehicle treatment. (*D*) Fractions collected every 10 minutes corresponded with the following conditions: 1-baseline, 2-baseline, 3-treatment, 4-treatment + 90 mM KCl, 5-wash. Prior treatment with compound ***45*** significantly reduced high potassium-evoked CGRP release in spinal cord (vs. DMSO, n=4-5, *P < 0.0001, one-way ANOVA with Dunnett’s post hoc comparisons).

### 3.5. 45 leads to accumulation of cytoplasmic CaV2.2 in the presence of proteasomal inhibition

A recent study demonstrated that the β subunit protects CaV2.2 from degradation by the proteasome [56]. To test if ***45*** could work through this pathway, DRG neurons were treated with ***45*** (20µM) or its vehicle (DMSO) for 1-16 hours and immunostained with an antibody against CaV2.2. Prolonged, but not short treatment, with ***45*** caused an increase in total CaV2.2 immunostaining (*Fig. S4*). To investigate a potential effect of ***45*** on CaV2.2 trafficking and degradation, DRG neurons were treated in presence or absence of the proteasome inhibitor lactacystin, and the ratio of membrane staining over cytoplasm staining was quantified as well as the total fluorescence intensity. DRG neurons cultured in presence of lactacystin exhibited an increase in total CaV2.2 staining (*Fig. S4C, E*), indicating that preventing protein degradation leads to increased levels of CaV2.2 expression. Levels of CaV2.2 in DRG neurons treated with ***45*** were not increased in presence of lactacystin (*Fig. S4*E). Taken together, these results raise the possibility that the increase in CaV2.2 expression levels after ***45*** treatment might be caused by a protective effect of ***45*** on CaV2.2 degradation; thus, diminishing the impact of proteasome inhibition on CaV2.2 expression levels. A higher portion of CaV2.2 channel appears to be located at the membrane in DRG neurons treated with ***45*** for 16 hours compared to neurons receiving both ***45*** and lactacystin (*Fig. S4F*). Blockade of the proteasome added to and ***45*** treatment might lead to an accumulation of CaV2.2 in the cytoplasm.

### 3.6. Inhibition of spinal CGRP release by 45

CaV2.2 activity controls C-fiber sEPSCs and the release of the nociceptive neurotransmitter calcitonin-gene related peptide (CGRP). Since ***45*** inhibited CaV2.2 and sEPSC frequency, we hypothesized that evoked spinal CGRP release would be inhibited by ***45***. To test this, we measured CGRP release evoked by depolarization (90 mM KCl, a concentration activating mostly CaV2.x channels [58]) in an ex-vivo preparation of the lumbar region of rat spinal cord. Enzyme-linked immunosorbent assay (ELISA) was used to measure CGRP content; samples were collected every 10 min. Basal CGRP levels were ~8.2 pg.ml^-1^.mg^-1^ of tissue (*Fig. 4D*, fractions #1 & 2). Prior to stimulation, control (0.1% DMSO) or a 20 μM ***45*** was added (*Fig. 4D*, fraction #3). While ***45*** application did not elicit any CGRP release from the spinal cords (*Fig. 4D*, fraction #3), depolarization evoked CGRP release from spinal cord which was inhibited by ~63 % by compound ***45*** (*Fig. 4D*, fraction #4). These results show that inhibiting CaV2.2 by targeting the CaVβ-CaVα interface with ***45*** results in decreased depolarization-evoked CGRP release.

### 3.7. Broad antinociceptive efficacy of 45

The peptide ω-conotoxin GVIA remains a defining ligand for CaV2.2 and is marketed as Prialt® (Primary Alternative to Morphine) for relief of chronic and cancer-related pain [44], establishing the therapeutic value of targeting CaV2.2. ***45*** increased withdrawal latency to a heat stimulus in naïve rats (*Fig 5A*), demonstrating anti-nociceptive potential. Furthermore, CaV2.2 is known to be involved in post-surgical pain [17], neuropathic pain [62], chemotherapy induced neuropathy [24], HIV-induced sensory neuropathy [37] and Neurofibromatosis type 1 (NF1) related pain [40]. Thus, we performed an analgesic appraisal of ***45*** on heat and tactile sensitivity of rats *in vivo*. We chose to administer ***45*** intrathecally (2 µg in 5 µl) to directly reach the site of action of CaV2.2 in the dorsal horn; the pre-synaptic termini of pain fibers that synapse with spinal cord neurons and require CaV2.2 for the release of neurotransmitter at these synapses. ***45*** reversed mechanical allodynia and thermal hyperalgesia in models of NF1-related pain (*Fig. 5B*), post-surgical pain (*Fig. 5C-D*) and neuropathic pain (*Fig. 5E-F*). ***45*** was also efficient in reversing mechanical allodynia in paclitaxel-induced peripheral neuropathy (*Fig. 5G*), and HIV-induced sensory neuropathy (*Fig. 5H*). The reversal lasted about 2-3 hours across the behavioral paradigms tested, consistent with a plasma half-life of 2.2 hr for ***45*** (data not shown). These results demonstrate the utility of ***45*** to modulate painful sensations in rodents.

**Figure 5.**
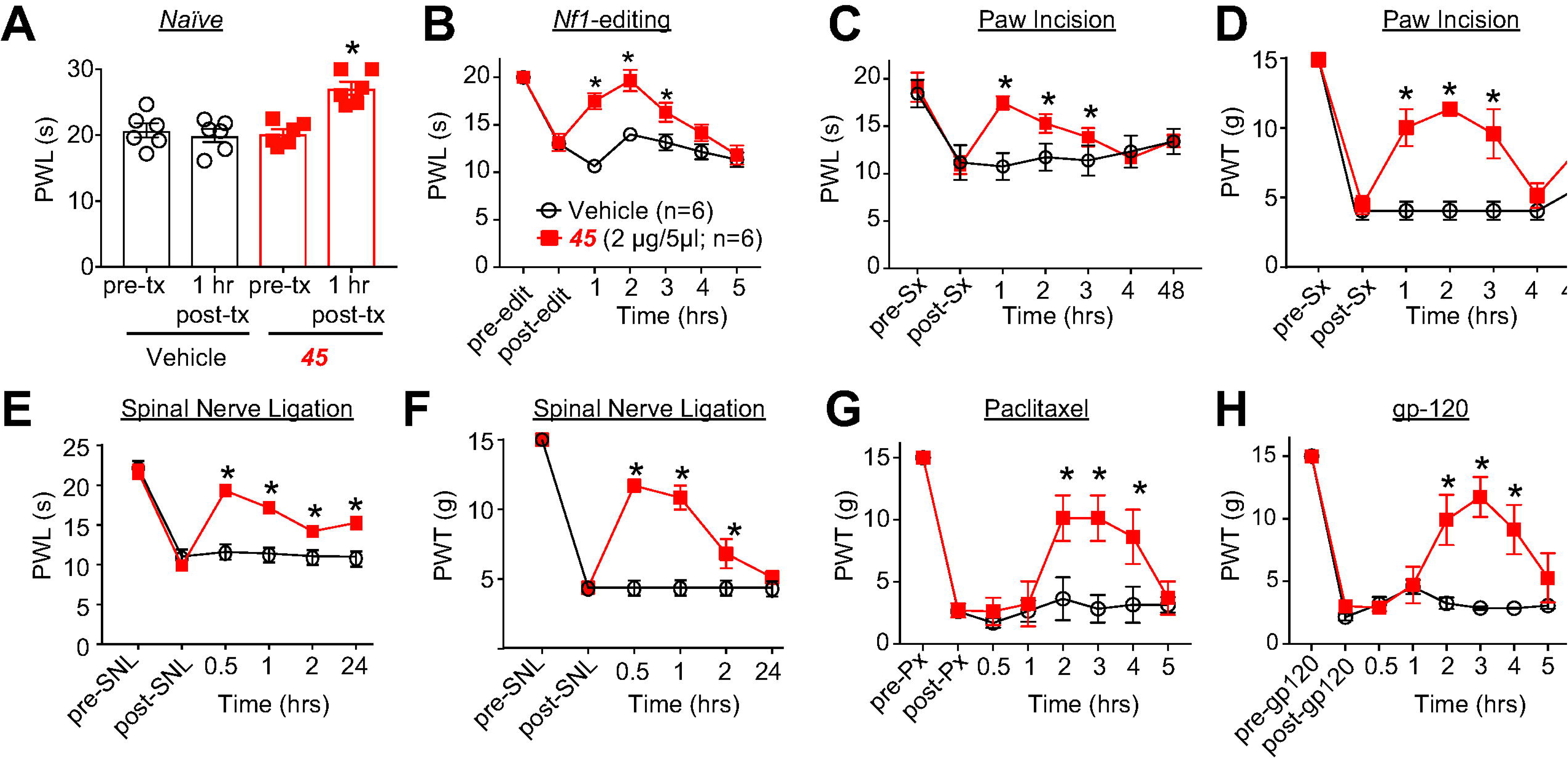
*45* is antinociceptive in naïve animals and reverses mechanical allodynia and thermal hyperalgesia in acute and neuropathic pain models. (*A*) Paw withdrawal latencies (PWLs) of naïve rodents to a heat stimulus were significantly increased 1 hour following an intrathecal (i.t.) injection of ***45*** compared to animals injected with vehicle (10% DMSO, 10% Tween80, 80% saline) (compared to i.t. injection of vehicle (n=6, *P < 0.05, two-way ANOVA). (*B*) PWLs of rodents subjected to targeted Cas9-mediated editing of *Nf1* (via i.t. injection) were significantly decreased, but i.t. injections of ***45*** reversed this behavior (vs. i.t. injection of vehicle, n=6, *P < 0.05, two-way ANOVA). (*C*) PWLs of rodents that received a paw incision (Sx) on the left hind paw were significantly decreased. I.t. injection of ***45*** reversed this nociceptive behavior (compared to i.t. injection of vehicle, n=6, *P < 0.05, two-way ANOVA). (*D*) Paw withdrawal thresholds (PWTs) of rodents that received a paw incision (Sx) on the left hind paw were significantly decreased. Intrathecal (i.t.) injection of ***45*** reversed this nociceptive behavior (compared to i.t. injection of vehicle, n=6, *P < 0.05, two-way ANOVA). (*E*) PWLs of rodents that received a spinal nerve ligation (SNL) on the left hind paw were significantly decreased. Intrathecal (i.t.) injection of ***45*** reversed this nociceptive behavior (compared to i.t. injection of vehicle, n=6, *P < 0.05, two-way ANOVA). (*F*) PWTs of SNL-rodents were also significantly decreased. I.t. injection of ***45*** also reversed this behavior (vs. i.t. injection of vehicle, n=6, *P < 0.05, two-way ANOVA). (*G*) PWTs of rodents that received four 2 mg/kg intraperitoneal (i.p.) injections of Paclitaxel (Px) over 12 days were significantly decreased. I.t. injection of ***45*** reversed this behavior (vs. i.t. injection of vehicle, n=6, *P < 0.05, two-way ANOVA). (*H*) PWTs of rodents were significantly decreased 15 days after receiving three i.t. injections of gp-120. I.t. injection of ***45*** reversed this behavior (vs. i.t. injection of vehicle, n=6, *P < 0.05, two-way ANOVA). The experiments were conducted in a blinded fashion.

### 3.8. 45 does not cause motor impairment

The clinically used CaV2.2 drug Prialt (Zinocotide) produces akinesia when injected into the brain [34]. To test if ***45*** would elicit a similarly undesirable effect, we used the rotarod test to assess rats’ motor function after an intracerebroventricular (i.c.v.) injection of ***45*** (2 µg in 5 µl). Rats were followed for 90 min after the injection and no motor impairment was observed (*Fig. S5*) after ***45*** brain injection. We did not observe other behavioral alterations such as seizures in any of the animals tested.

### 3.9. 45 works through a spinal mechanism and does not affect anxiety

***45*** was tested in acute thermal pain models (hot plate or tail-flick) as CaV2.2 blockers have been reported to be more effective against chronic than acute pain [33]. When a dose (15 mg/kg) of ***45*** was intraperitoneally administered in acute thermal pain models, the latency times were not affected in the hot plate test (predominantly supraspinal) but increased in the tail-flick test (predominantly spinal) (*Fig. 6A, B*).

**Figure 6.**
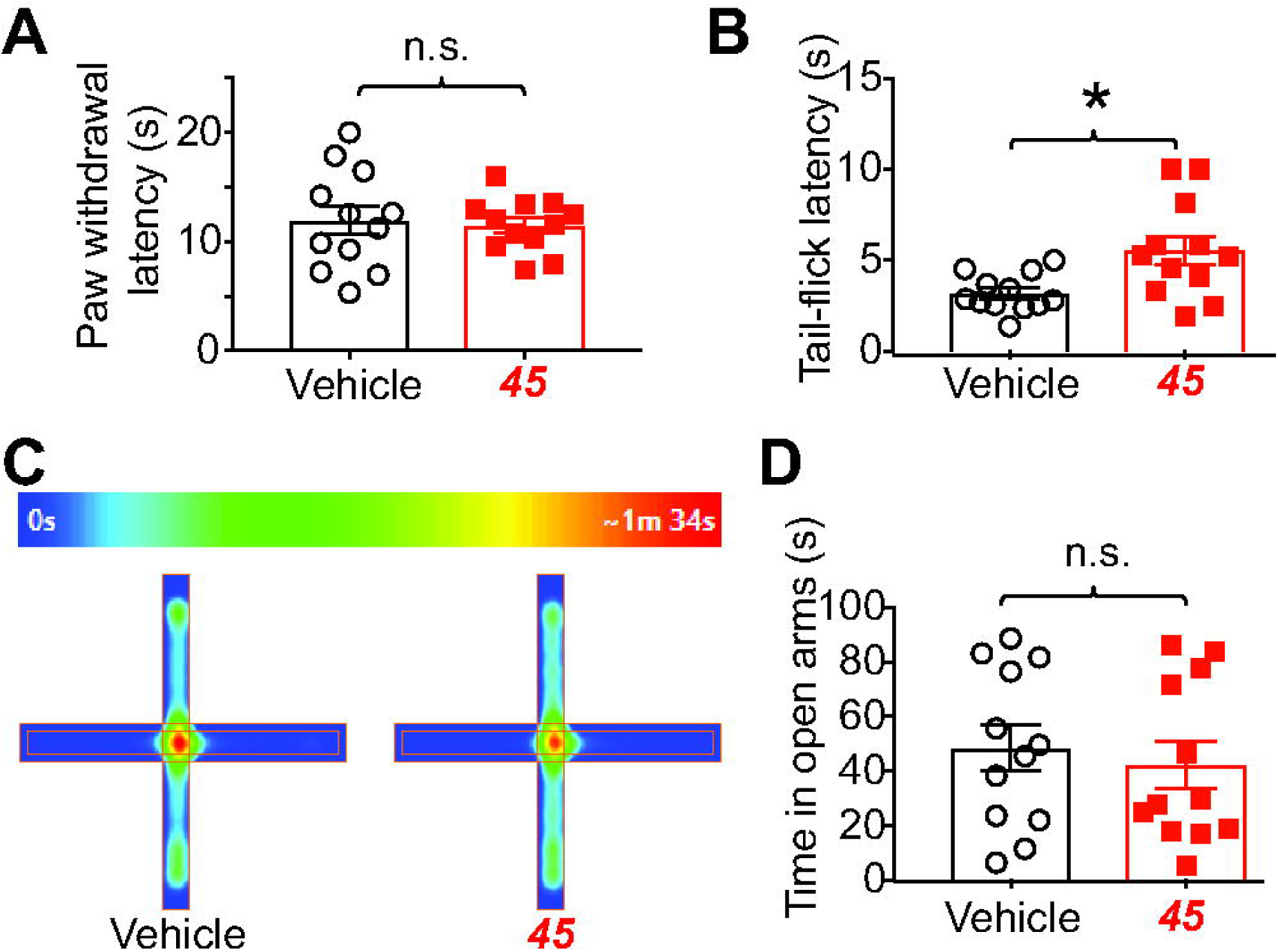
*45* is analgesic in mice but does not affect anxiety-related behaviors. (*A*) Paw withdrawal latencies of naïve mice to a hot-plate test (52°C) 75 min after injection of ***45*** administered intraperitoneally (i.p.) were unaffected compared to animals injected with vehicle (10% DMSO, 10% Tween80, 80% saline) (compared to i.p. injection of vehicle (n=12, p=0.7583, t-test with Welch’s correction)). (*B*) Tail-flick latencies of naïve mice to a hot (52°C) water bath 75 min after injection of ***45*** administered intraperitoneally were significantly increased compared to animals injected with vehicle (n=12, *P < 0.05, t-test with Welch’s correction). Naïve mice received an i.p. injection of ***45*** (15mg/kg) or its vehicle and anxiety-related behaviors were assessed during 10 minutes in an elevated plus maze test. (*C*) Heatmaps of the positions occupied by mice treated with in the elevated-plus maze apparatus. (*D*) Bar graphs of the mean time spent in the open arms of the elevated plus maze. No differences were found between animals treated with ***45*** or with its vehicle (p=0.6014, t-test with Welch’s correction, n=12 mice/group). The experiments were conducted in a blinded fashion.

As it has been previously demonstrated that inhibition of CaV2.2-mediated signaling induces alterations in the neuronal network involved in anxiety-related behaviors [29; 64], we next tested the effects of ***45*** on anxiety using the elevated plus maze test. Time spent in the open arms of the elevated plus maze, time spent in the closed arms and number of entries in the open arms did not differ between animal treated with ***45*** or its vehicle (*Fig. 6C, D*), suggesting that ***45*** does not affect anxiety-related behaviors.

## 4. Discussion

The present work used a rational structure-based design strategy to identify a new class of non-opioid, analgesic compounds targeting the CaVα/CaVβ interface. The quinazoline analog ***45*** (i) specifically bound to CaVβ, (ii) inhibited the biochemical interaction between CaVβ2 and the AID of CaV2.2 and to a lesser extent between CaVβ2 and the AID of CaV2.1, but not other subtypes, (iii) and functionally inhibited CaV2.2 function in DRG sensory neurons. ***45*** inhibited the pre-synaptic localization of CaV2.2 *in vivo* as well as inhibiting spinal neurotransmission, which resulted in decreased neurotransmitter release from the spinal cord. Finally, ***45*** was anti-nociceptive in naïve rats and reversed mechanical allodynia and thermal hyperalgesia in rodent models of acute (post-surgical), neuropathic (spared nerve ligation, paclitaxel or HIV-induced sensory neuropathy) and genetic (NF1-related) pain. No adverse effects were observed for ***45*** in rats or mice. These results are an instructive example that pharmacological disruption of the CaVα/CaVβ interface can result in a selective, safe and broadly antinociceptive compound, setting the stage for structure-activity relationship studies to improve on this lead molecule for the development of novel pain therapeutics.

The CaVα/CaVβ interaction shapes activity and trafficking of all subtypes of high voltage-gated calcium channels (i.e., L-, N-, P/Q-, and R-type). Prior to the crystallization of CaVβ subunits, Wyeth-Ayerst Research used a high throughput yeast two-hybrid screen to identity compounds that disrupt α1b and β3 and found WAY141520, which inhibited calcium currents with an IC50 of 95 μM with some inhibitory activity against CaV2.2 but was not tested against other channels nor was its mechanism of action studied [63]. Structural studies of CaVβs in early 2000’s [11; 42; 52] with a fragment of the α subunit revealed a high homology between the various α-β subunits, predicting that targeting of this interface would result in non-selective compounds. The details of this interface have been extensively studied in deep thermodynamic detail for all of the CaV1 and CaV2 AIDs and that has defined the hotspots on both sides of this interaction [53]. Of the three “hot spots” identified by the Minor laboratory, V241 makes no contacts with the AID, N390 does not contribute to the energetics of the interaction, and only I343 is important, interacting with one of the core residues that comprise the AID hotspot (CaV1.2 Y437) [53]. Despite these theoretical predictions, ***45*** exhibited selectivity for CaV2.2. ***45*** interacts with CaVα-CaVβ domain mainly at the lipophilic pocket defined by Val 339 – Ile 343 [52]. The AID sequence aligns (side chains of critical residues Y437, W440, and I441; *Fig. 1C*) with the quinazoline ring and phenylpropyl group of ***45***. We hypothesize that these interactions result in high selectivity for ***45***. That ***45*** reduced the binding affinity between CaVβ and AID peptides of CaV2.2 (by ~17.3-fold) and CaV2.1 (by ~5-fold) but had no effect on binding to other channel subtypes demonstrates selectivity for the interaction. Although this selectivity was unexpected, the electrophysiology recordings on sensory neurons confirmed a selectivity for CaV2.2 while the electrophysiology recordings in heterologous cells, performed in an independent laboratory, confirmed inhibition through CaV2.2 and CaVβ1b, CaVβ2a, and CaVβ3. Inhibiting CaV2.2 with ***45*** did not show the typical regulation of the channel’s gating property by CaVβs [8]. But this is consistent with a prior report that demonstrated no effect on voltage-dependent activation of high-voltage activated calcium channels upon deletion of CaVβ3 subunit in DRGs [30]. It has also been reported that orientation of CaVβ to the α subunit of CaV2.2 subunit is critical for its regulation of channel activity with insertions or deletion of the AID linker, which are expected to maintain the α-helical structure of the linker but induce a rotation of CaVβ with respect to CaVα1, diminish Cavβ regulation of activation and inactivation [55]. Thus, ***45*** could possibly induce conformational changes that result in disrupt the rigidity of the α-helix which may in turn explain the lack of effects on biophysical properties.

That ***45*** was effective and safe *in vivo* likely stems from the restricted expression patterns of the CaVβ subunits. For example, CaVβ1 is expressed in skeletal muscle [21], CaVβ2 is expressed in the heart and at very low levels in the central nervous system (CNS) [31], CaVβ3 expression is found in smooth muscle and in the CNS (including the spinal cord) [31], and CaVβ4 is expressed predominantly in the cerebellum [31]. CaV2.2 is expressed in the brain and in the sensory neurons [10]. Thus, only in the brain and sensory neurons does CaV2.2 expression overlap with that of CaVβ3 and CaVβ4. Our *in vitro* results demonstrated that only coexpression of the CaVβ3/4 subunits with CaV2.2 resulted in high Ca^2+^ currents while CaVβ1/2 coexpression with CaV2.2 (a configuration that is not physiologically relevant because these subunits are primarily expressed in skeletal or heart muscles) resulted in low Ca^2+^ currents. **45** inhibited Ca^2+^ currents when co-expressed with CaVβ1/2/3, but not CaVβ4. Thus *in vivo*, **45** would only target CaVβ3/CaV2.2 coupling because of the overlapping tissue distribution of these subunits. The notion that CaV2.2 functions with CaVβ3 to mediate nociception *in vivo* is supported by mouse models lacking either CaV2.2 [46] or CaVβ3 [41] with these transgenic mice demonstrating resistance to pain sensation. Notably, CaVβ3 expression is increased in neuropathic pain [30], in parallel with increased voltage-gated calcium channel function. Thus, the specific uncoupling of CaVβ3/CaV2.2, but not CaVβ4/CaV2.2, may underpin the calcium channel subtype selectivity of ***45*** and its *in vivo* efficacy for the reversal of chronic pain behaviors.

The time-course of action of ***45*** is reminiscent of Gabapentin, another trafficking regulator of the N-type channel, which requires at least 17 hours prior to inhibition of calcium currents with maximum inhibition observed at 40 hours [19]. From a mechanistic viewpoint, fractionation studies demonstrated that ***45*** reduces the pre-synaptic localization of CaV2.2 which may account for its effect on a reduction in neurotransmitter (CGRP) release. Long-term treatment of DRGs with ***45*** increased CaV2.2 expression either due to increased protein synthesis or to a reduction of its degradation. Consistent with the work of Dolphin and colleagues [56], our data suggests that ***45*** may act in a fashion analogous to the β subunit in sparing the channel from proteasomal degradation.

A common denominator in the neuropathic models appraised here is the plasticity of CaV2.2: targeted truncation of neurofibromin, achieved by acute CRISPR/Cas9 editing of the *Nf1* gene in adult rats, resulted in increases of CaV2.2 currents [39]; SNL increased CaV2.2 expression in the synaptic plasma membranes of the dorsal horn [28]; and the *Cacna1b* gene encoding CaV2.2 is highly upregulated in a rodent model of HIV-SN treated with gp120 compared to sham-treated controls [32]. Previous reports have shown attenuation of allodynia by a CaV2.2-blocking compound in the paw incision model of surgical pain [17] as well as the paclitaxel-induced model of chemotherapy-induced peripheral neuropathy [5]. The broad antinociceptive potential of ***45*** across acute and neuropathic models, in two species, can be attributed to its biochemical and functional inhibition of the CaVα-CaVβ interaction, a mechanism distinct from direct block of CaV2.2 by small molecules covering various pharmacophores [61].

Therapeutic management strategies for chronic pain usually involve the use of opioid molecules. These compounds primarily act on the MOR expressed on DRG sensory neurons to provide analgesia. MOR inhibition of nociception is mediated by the activation of G(i/o)-type GTP binding protein which inhibits CaV2.2 [35; 36]. In neuropathic pain, alternative splicing of CaV2.2 mRNA leads to two different CaV2.2 isoforms containing either the exon 37a (e37a) or e37b in sensory neurons [4]. After a neuropathic injury, expression of e37a is lost, resulting in an enrichment of e37b containing CaV2.2 in sensory neurons [1]. This switch in alternative splicing is an important event underlying resistance and tolerance to opioids since the remaining e37b containing CaV2.2 is resistant to the inhibition mediated by opioids [2]. No alternative splicing has been reported in the binding region between CaVα/CaVβ. Thus, targeting this interface using ***45*** is unlikely to be susceptible to similar tolerance mechanisms in neuropathic pain.

Although the CaV2.2 inhibiting drug Ziconotide (Prialt®) is effective in chronic and cancer-related pain [44], it requires intrathecal dosing to circumvent systemic side effects that include confusion, depression, hallucinations, decreased alertness, somnolence, orthostatic hypotension and nausea [48]. Thus, a small molecule, such as ***45***, devoid of these encumbrances and lacking a non-opioid mechanism would be a welcome addition to the armamentarium of the pain physician for managing chronic pain.

## Supporting information

## Acknowledgements

This work was supported by a Career Development Award from the Arizona Health Science Center to M.K., grants from the National Natural Science Foundation of China (81603088) to J.Y., National Institutes of Health awards (NINDS K08 to A.P.; R01NS098772 from the National Institute of Neurological Disorders and Stroke and R01DA042852 from the National Institute on Drug Abuse to R.K.); and a Neurofibromatosis New Investigator Award from the Department of Defense Congressionally Directed Military Medical Research and Development Program (NF1000099) to R.K.

