## Supplementary material for "Targeting the CaVα-β interaction yields a selective antagonist of the N-type CaV2.2 channel with broad antinociceptive efficacy"

[1]Department of Pharmacology, College of Medicine, University of Arizona; [2]The BIO5 Institute University of Arizona; [3]Department of Physiology and Cellular Biophysics, and Pharmacology, Columbia University College of Physicians and Surgeons; [4]Convergence Research Center for Dementia, Korea Institute of Science and Technology, Seoul 02792, Korea; [5]Division of Bio-Medical Science &Technology, KIST School, Korea University of Science and Technology, Seoul 02792, Korea; [6]KHU-KIST Department of Converging Science and Technology, Kyung Hee University, Seoul 02447, Korea; [7]Department of Anesthesiology, College of Medicine, University of Arizona; and [8]The Center for Innovation in Brain Sciences, The University of Arizona Health Sciences, Tucson, Arizona; and

[a]co-first authors

**Corresponding Author*: Dr. Rajesh Khanna, Department of Pharmacology, College of Medicine, University of Arizona, 1501 North Campbell Drive, P.O. Box 245050, Tucson, AZ 85724, USA Office phone: (520) 626-4281; Fax: (520) 626-2204;

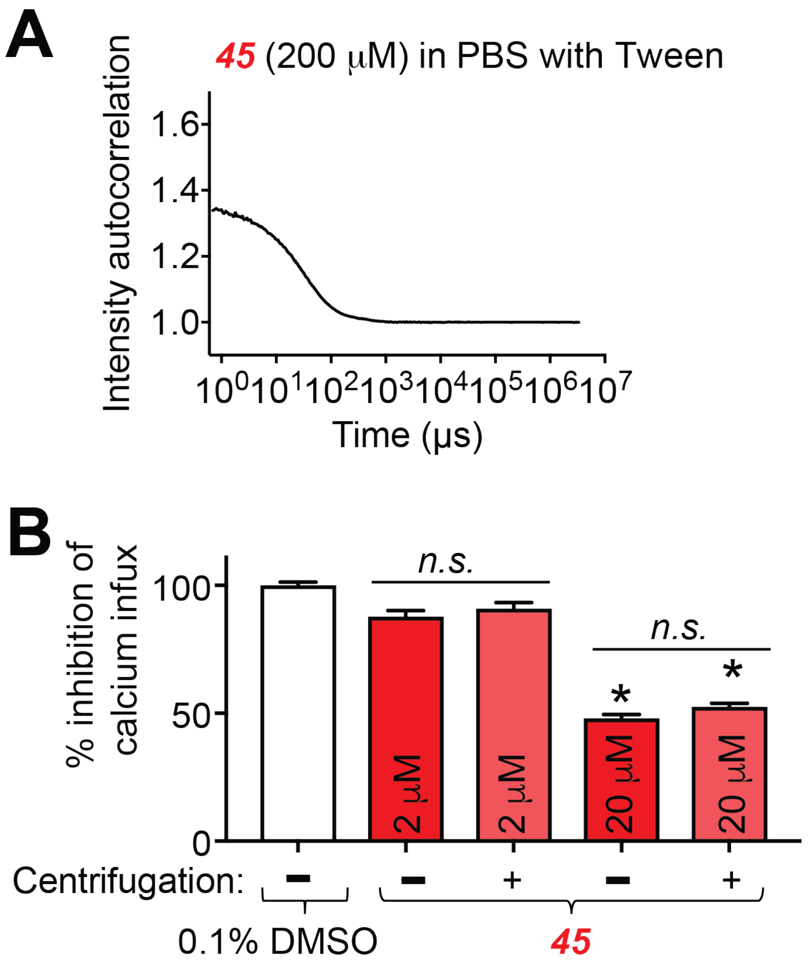


**Figure S1. *45* does not aggregate at relevant concentrations nor does it suffer from any loss of ability to inhibit calcium influx when subjected to ultracentrifugation.** (A) Querying ***45*** in the Aggregator Advisor database (http://advisor.bkslab.org), developed by Dr. Brian K. Shoichet's laboratory at UCSF, with the SMILES ID (Cc1noc(C)c1CC(=O)NCc4nc(NCCCc2ccccc2)c3ccccc3n4) revealed no similarity to known aggregators. A similar query in the database Zinc15 http://zinc15.docking.org/patterns/home/ revealed no hits to compounds matching PAINS or similar to known aggregators. Since the Aggregator Advisor calculated a cLogP value of 3.5, which is in the range reported for many other aggregators, dynamic light scattering was performed. Dynamic light scattering curve of ***45*** in the presence of a non-ionic detergent (0.1% Tween-20) did not reveal significant colloidal aggregation with less than 9% of the compound forming particles around a 50 nm radius. (B) To control for potential effects of aggregation, we subjected ***45*** to a centrifugation spin-down (15 min at 21,000 X g) and then used the supernatant from the spin-down of ***45*** to perform calcium imaging studies.
Bar graphs show normalized peak calcium response average ± S.E.M. of DRG sensory neurons incubated overnight with a 2 or 20 μM concentration of ***45*** and vehicle in response to 90mM KCl. Responses were normalized to that of the DMSO. Asterisks indicate statistical significance compared with cells treated with the DMSO vehicle ($p<0.05$, one-way ANOVA with Dunnett's post-hoc test). There were no differences (not significant, *n.s.*) in the responses between cells treated with ***45*** irrespective of whether it was centrifuged or not.

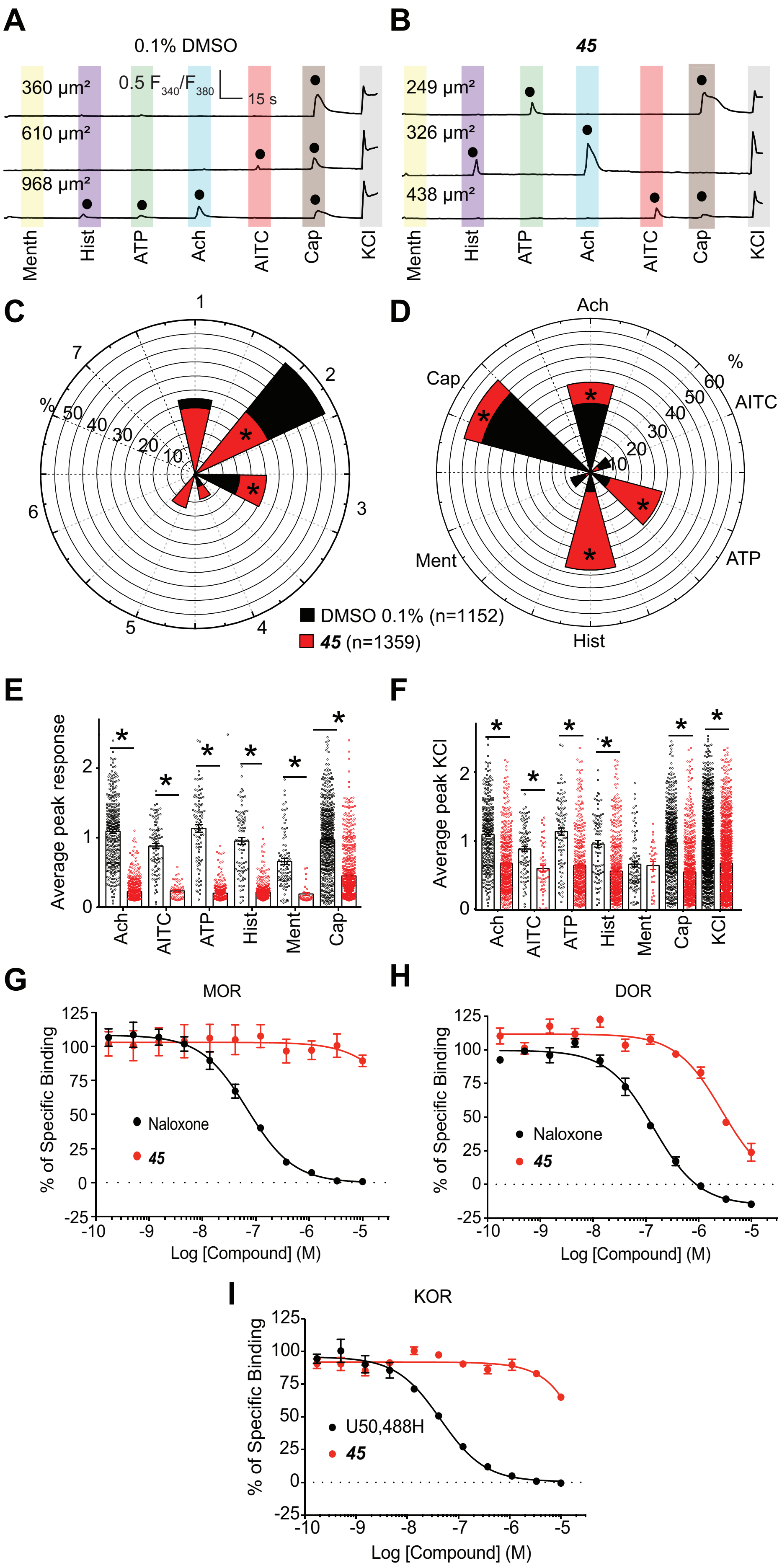
A
0.1% DMSO
360 µm²
610 µm²
968 µm²
0.5 F340/F380
15 s
Menth
Hist
ATP
Ach
AITC
Cap
KCl
B
45
249 µm²
326 µm²
438 µm²
Menth
Hist
ATP
Ach
AITC
Cap
KCl
C
1
2
3
4
5
6
7
%
50
40
30
20
10
D
Ach
AITC
ATP
Hist
Ment
Cap
%
60
50
40
30
20
10
DMSO 0.1% (n=1152)
45 (n=1359)
E
Average peak response
Ach
AITC
ATP
Hist
Ment
Cap
F
Average peak KCl
Ach
AITC
ATP
Hist
Ment
Cap
KCl
G
MOR
% of Specific Binding
Naloxone
45
Log [Compound] (M)
H
DOR
% of Specific Binding
Naloxone
45
Log [Compound] (M)
I
KOR
% of Specific Binding
U50,488H
45
Log [Compound] (M)

**Figure S2 (see previous page). Identifying potential off-target effects of compound *45* among pain-associated channels and receptors.** (*A*) Representative $Ca^{2+}$ imaging traces from 0.1% DMSO (n=1152) and (*B*) 30μM compound ***45*** (n=1359) treated DRGs. This data includes DRGs analyzed over two independent experiments with up to nine trials per experiment. During each trial, >100 DRGs were sequentially stimulated for ~15 seconds (indicated by colored bars) with menthol (400 nM), histamine (50 μM), adenosine triphosphate (ATP) (10 μM), allyl isothiocyanate (AITC) (200 μM), acetylcholine (1 mM), capsaicin (100 nM), and KCl (90 mM). Only DRGs responsive to membrane depolarization-evoked $Ca^{2+}$ influx by KCl were included. Furthermore, only responses greater than 10% of baseline were included (indicated by filled black circles). The size of each cell is indicated by the surface area value in each trace. (*C*) Analysis of DRG responses revealed the percentage of cells responding to the indicated number of molecular agonists, independent of which agonists elicited a response, in both DMSO and ***45*** treated conditions (n=184-588, *P < 0.001, z-test.) (*D*) Analysis of DRG responses revealed the percentage of cells responding to the indicated molecular agonist, independent of any other agonist that also elicited a response, in both DMSO and ***45*** treated DRGs (n=88-693, *P < 0.01, z-test). (*E*) Summary bar graph shows the average peak response to each molecular agonist in DRGs treated with DMSO or compound ***45*** (n=1152-1359, *P < 0.05, Student's *t* test). (*F*) Summary bar graph shows the average peak KCl response among DRG subclasses responsive to the indicated molecular agonist following treatment with DMSO or ***45*** (n=1152-1359, *P < 0.05, one-way ANOVA). (*G*) Competition radioligand binding was performed in CHO cells expressing the human MOR, DOR, or KOR (see Methods for details). Compound ***45*** or a positive control compound competed against $^3H$-diprenorphine in all 3 cell lines. Curves reported as the mean ± SEM of the mean value from each individual experiment in N = 3 independent experiments. The $K_I$ also reported as the mean ± SEM of the individual value from each of N = 3 independent experiments. Compound *45* did not bind to the MOR. Naloxone $K_I$ = 33.9 ± 1.7 nM. (*H*) Compound ***45*** produced 76.2% inhibition at 10 μM at the DOR, with an incomplete curve. This indicates an $IC_{50}$ > 3.3 μM and a $K_I$ > 1 μM. Naloxone $K_I$ = 55.7 ± 6.7 nM. (*I*) Compound ***45*** did not appreciably bind to the KOR. U50,488 $K_I$ = 22.4 ± 4.1 nM.

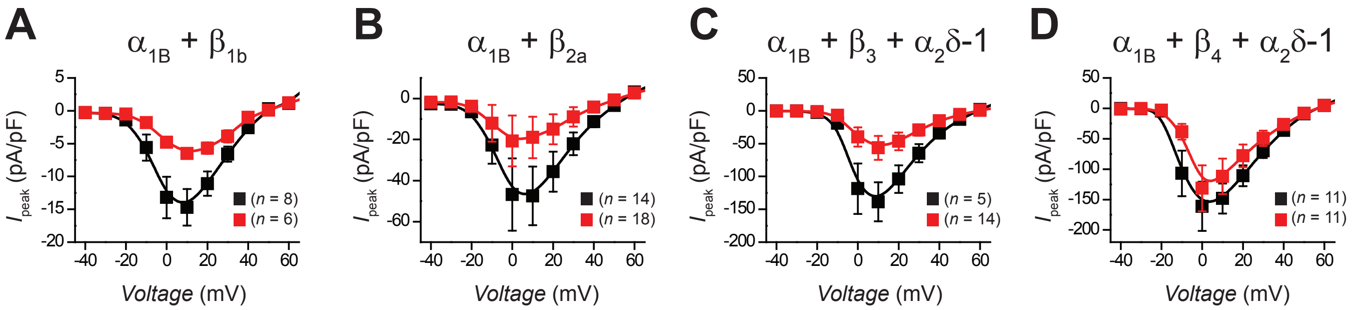


**Figure S3. *45* disrupts α1B-β1b, α1B-β2a, and α1B-β3-α2δ-1 interactions to inhibit $Ca^{2+}$ currents.** HEK293 cells expressing either (A) α1B+β1b, (B) α1B+β2a, (C) α1B+β3+α2δ-1, or (D) α1B+β4+α2δ-1 were subjected to an activation voltage step protocol to determine their I-V relationship following overnight treatment with 0.1% DMSO (black) or 30μM compound ***45*** (red). $Ca^{2+}$ currents in α1B+β1b, α1B+β2a, and α1B+β3+α2δ-1 expressing cells were significantly reduced in compound ***45*** treated DRGs compared to 0.1% DMSO treated counterparts (n=5- 18, *P<0.05, Kruskal-Wallis test with Dunnett's post hoc comparisons).

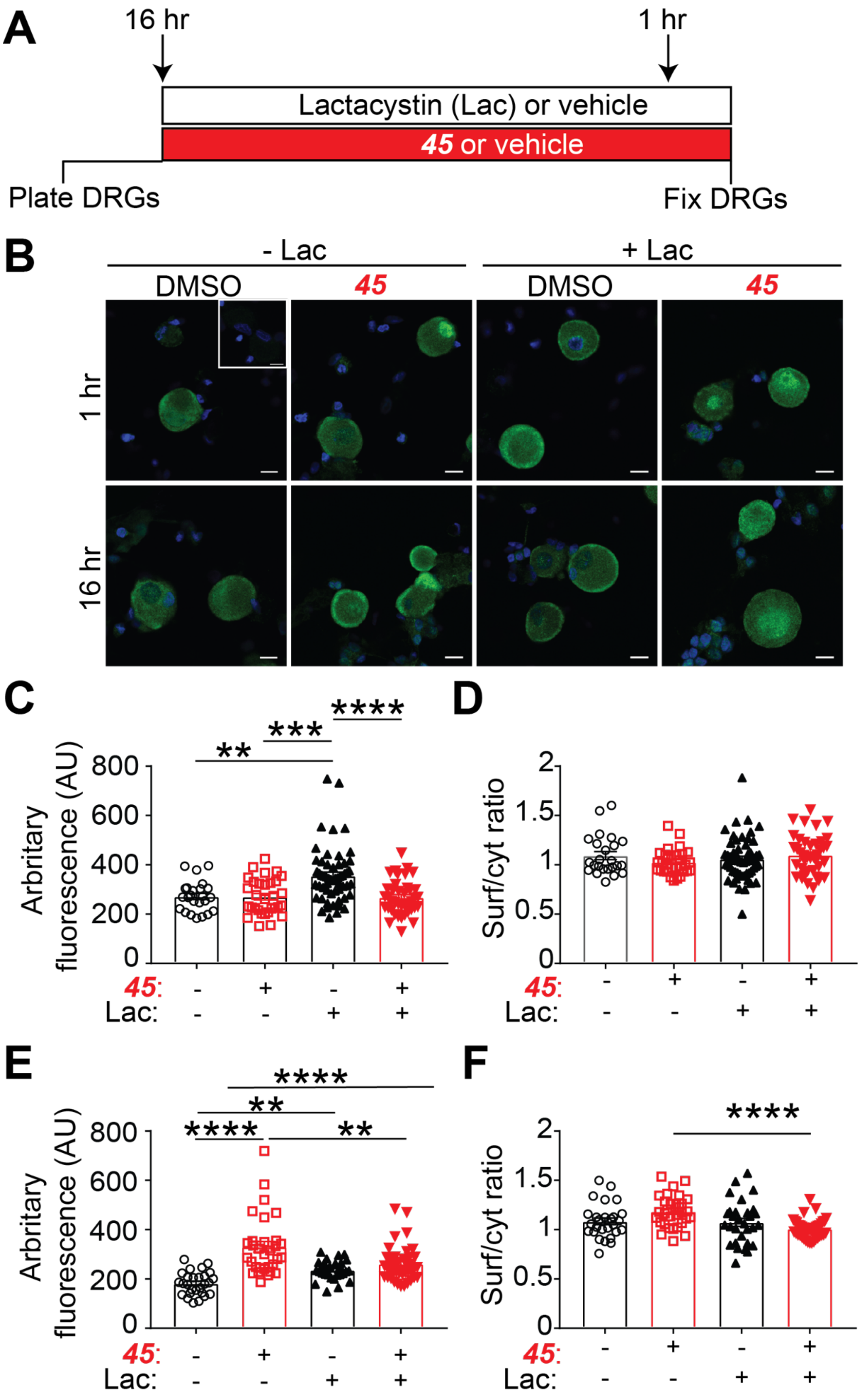


**Figure S4. Blockade of the proteasome and *45* treatment leads to an accumulation of CaV2.2 in the cytoplasm.** (A) Timeline of DRG neurons treatment with ***45*** (20μM) or its vehicle (DMSO) and lactacystin (10μM) and representative micrographs of DRG neurons immunostained with CaV2.2. Nuclei were counterstained with DAPI. For the negative control (insert), primary antibody was omitted.

Scale bars: 10µm. (B) Bar graphs of mean fluorescence intensity for each cell and ratio of membrane fluorescence over cytoplasmic fluorescence (Surf/Cyt ratio) for each cell after a 1-hour treatment with ***45*** or DMSO. (C) Bar graphs of mean fluorescence intensity for each cell and ratio of membrane fluorescence over cytoplasmic fluorescence for each cell after a 16-hours treatment with ***45*** or DMSO. n=24 to 56 cells/condition. *$p<0.05$, **$p<0.01$, ***$p<0.005$, ****$p<0.001$ Kruskall-Wallis tests followed by Dunnett's multiple comparisons tests.

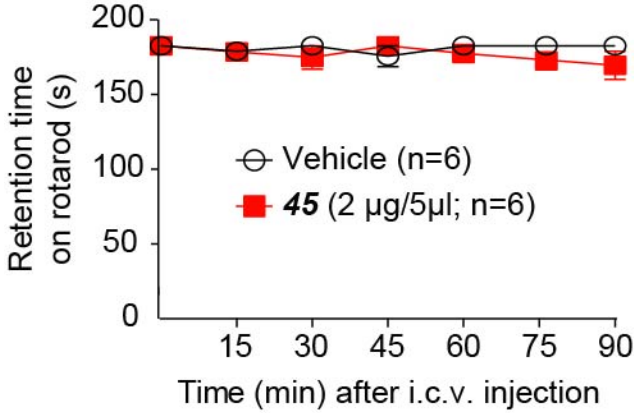


**Figure S5. Compound *45* does not impair locomotion.** Rodents were subjected to intracerebroventricular (i.c.v.) injection of ***45*** or vehicle. In a rotarod test, no significant difference in performance (measured by retention time) was observed between the two cohorts ($p>0.05$).

### Supplementary Methods

**Reagents.** All chemicals, unless noted, were purchased from Sigma (St. Louis, MO).

**Animals.** Pathogen-free, adult male Sprague-Dawley rats (225- 250 g; Envigo, Indianapolis, IN) were housed in temperature (23 ± 3°C) and light (12-hour light/12-hour dark cycle; lights on 07:00-19:00) controlled rooms with standard rodent chow and water available ad libitum. The Institutional Animal Care and Use Committee of the College of Medicine at the University of Arizona approved all experiments. All procedures were conducted in accordance with the Guide for Care and Use of Laboratory Animals published by the National Institutes of Health and the ethical guidelines of the International Association for the Study of Pain. Experimenters were blinded to the experimental groups and treatments to which the animals were randomly assigned.

**In silico docking and software for structural representation.** Virtual screening workflow was done in Schrodinger – Glide molecular modeling software using the X-ray crystal structure of the complex between CaVβ2a subunit and a peptide of the α1c subunit (PDB code:1T0J) [29]. The α1c subunit peptide was removed and its binding site was used for docking of 50,000 drug-like small molecule library (molecular weight ≤ 500 Da) available from ChemBridge Inc. The resulting complexes were ranked using Glide XP score and other energy related terms. Forty-nine compounds were analyzed for their binding in the α-binding pocket on CaVβ2a. Selected compound were purchased and tested for activity.

**Dynamic Light Scattering.** Particle formation was measured using a DynaPro MS/X (Wyatt Technology) as previously described by the Shoichet and colleagues[4]. ***45*** was measured both in the absence and in the presence of 0.1% Tween-20.

**Protein purification.** Codon optimized sequence (E. coli) for CaVβ2a residues 203-425 was inserted between the restriction sites EcoRI-SalI in pET-28a(+) plasmid (synthesized by Genscript). BL21 cells expressing CaVβ2a were resuspended in 50 mM sodium phosphate pH7.5, 500 mM NaCl, 10% glycerol, supplemented with Complete EDTA-free protease inhibitors (Roche, Basel, Switzerland). Disruption of the bacteria was performed by two rounds of high-pressure homogenization at 10,000 PSI with a LM10 microfluidizer (Microfluidics, Westwood), and the lysate was centrifuged 45 min at 4,500xg at 4°C. The supernatant was loaded on a His-Trap column (GE Healthcare, Uppsala, Sweden) equilibrated with 50 mM HEPES pH7.5, 10 mM imidazole, 500 mM NaCl, 10% glycerol, 0.5mM DTT. After a washing step with 50 mM HEPES pH7.5, 50 mM imidazole, 500 mM NaCl, 10% glycerol, 0.5mM DTT, CaVβ2a was eluted with a gradient of imidazole. The fractions of interest were loaded on a HiLoad Superdex size exclusion column (GE Healthcare, Uppsala, Sweden) and eluted with a gradient of imidazole. Protein concentration was determined by a Pierce assay using bovine serum albumin as a standard. The purity of the protein was verified with SDS-PAGE. The proteins were flash frozen in liquid nitrogen and stored at −80°C.

**Saturation Transfer Difference Nuclear Magnetic Resonance Spectroscopy.** 1D $^1$H STD NMR [14] for ***45*** (1 mM in 100 mM sodium phosphate pH 6.9, 10 % $D_2O$) in the absence of protein was acquired at 25 °C on a Varian Inova 600 MHz spectrometer equipped with a Varian cold TR/PFG probe. 1D $^1$H saturation transfer difference nuclear magnetic resonance (STD NMR) [13] spectra with a spectral width of 12 ppm were collected for samples containing 100 μM ***45*** and CaVβ2a in phosphate buffer. STD NMR spectra were collected with a spectral width of 12 ppm, 16 K data points, and 3 s repetition delay. A saturation of the protein was achieved by a 2-s train of selective 50 ms Gaussian pulses centered at 0.74 ppm (on resonance) and 30 ppm (off resonance). A 20-ms spin-lock was used to suppress the protein signal, followed by the double PFG spin echo to remove residual water signal. We acquired 512 scans per experiment. The on-resonance and off-resonance spectra were acquired interleaved, and the difference spectrum was acquired by phase cycling. Spectra processing and analysis were performed with the VNMRJ 3.2 and MestReNova 7.1.

**Microscale Thermophoresis.** MST, a method that monitors the thermophoretic movement of molecules in optically generated microscopic temperature gradients thereby permitting analysis of biomolecular interaction was performed as described previously [30; 31]. In MST, increasing concentrations of unlabeled ligand are mixed with the fluorescently labeled biomolecule, kept at constant concentration. Purified Beta2-CaV-His was fluorescently labelled using the His-Tag labeling kit RED-Tris-NTA 2$^{nd}$ generation (Nanotemper, Germany) according to the manufacturer's instructions. Briefly, Beta2-CaV-His was diluted to 200 nM in PBS supplemented with 0.1% Tween-20 (PBS-T buffer) and 0.1% PEG 8,000 and mixed with one molar equivalent of the fluorescent NT-647-His-labeling dye and incubated for 30 min at room temperature. Labeled Beta2-CaV was then centrifuged at 15,000Xg for 10 min at 4°C. 25 nM of labeled Beta2-CaV-His was mixed with increasing concentrations of AID-CaV2.2 peptide in PBS-T, 0.1% PEG 8,000 buffer and incubated 10 min at room temperature. The thermophoresis measurements were performed on a Monolith NT.115 (Nanotemper, Germany) using MST premium capillaries, at 100% LED at high MST power. Data analysis was performed with the MO Affinity Analysis software (Nanotemper) using the Kd model (standard fitting model derived from law of mass action).

**MOR-CHO, DOR-CHO, and KOR-CHO Cell Lines and Cell Culture.** The MOR-CHO cell line was purchased from PerkinElmer (#ES-542-C). The DOR cell line was created and characterized in our lab, as previously reported [26]. The KOR cell line was also created in our lab; an N-terminal 3X-hemagglutinin tagged human KOR expression clone from Genecopoeia was electroporated into parental CHO cells and selected with 500 µg/mL G418. The resulting selected population was enriched for receptor expression by live cell labeling with anti-HA-Alexa488 antibody and separating the top 2% of the population by flow cytometry. This high expressing population was characterized by immunocytochemistry and Western blot to establish receptor expression and expected signal transduction activation. All 3 cell lines were characterized by saturation radioligand binding with $^{3}$H-diprenorphine, and the measured $K_D$ used in competition binding experiments to calculate the $K_I$ (MOR = 5.23 nM; DOR = 0.93 nM; KOR = 1.81 nM; all the mean of N ≥ 3 independent experiments). All cells were cultured in 50:50 DMEM/F12 media with 10% heat-inactivated FBS and 1X penicillin/streptomycin supplement (all Gibco/ThermoFisher brand) in a 37°C humidified incubator with 5% $CO_2$ atmosphere; propagation cultures were further maintained in 500 μg/mL G418. Cultures were propagated for no more than 20 passages before discarding. Cell pellets for experiments were prepared by growth in 15 cm plates, harvest with 5 mM EDTA in dPBS (no calcium or magnesium), and stored at -80°C prior to use.

**HEK Cell Culture and Transfection.** Low-passage-number HEK (human embryonic kidney) 293 cells were transiently transfected with 8 μg of $\alpha_{1B}$-GFP, $\beta_{1b}$-[R/C/Y]FP, $\beta_{2a}$-[R/C/Y]FP, $\beta_3$-[R/C/Y]FP, $\beta_4$-[R/C/Y]FP, and/or $\alpha_2\delta$-1-[R/C/Y]FP, in addition to 3 μg of T antigen by calcium phosphate precipitation [5].

**Primary Cell Culture of DRGs.** Dorsal root ganglia from all levels were acutely dissociated using methods described previously [18; 20; 32]. Briefly, dorsal root ganglia (DRG) were removed from naive animals. The DRG were treated with collagenase, type I (5 mg/ml) and neutral protease (3.125 mg/ml) in bfDMEM for 45 minutes (Worthington Biochemical, Lakewood, NJ). The cells were then dissociated by mechanical trituration in culture media. The culture media was DMEM, supplemented with 10% fetal bovine serum, penicillin (100 mg/mL), streptomycin (100 U/mL), normocin (0.8 μg/ml) and nerve growth factor (30 ng/mL), Life Technologies, Carlsbad, CA). The cells were then plated (at 20 μl) on coverslips coated with poly-L-lysine and laminin (BD bioscience, San Jose, CA) and incubated for 1-2 hours before more culture media was added to the wells (up to 1 ml total volume). The cells were then allowed to sit undisturbed for 15 to 18 hours to adhere at 37°C (with 5% $CO_2$).

**Calcium imaging.** DRG neurons were loaded for 30 min at 37°C with 3 µM Fura-2 Low Affinity ($K_D$ = 0.14 µM, $\lambda_{ex}$ 340, 380 nm/$\lambda_{emi}$ 512 nm) to follow changes in intracellular calcium ($[Ca^{2+}]_c$) in a standard Tyrode's solution (at ~310 mOsm) containing 119 mM NaCl, 2.5 mM KCl, 2 mM $MgCl_2$, 2 mM $CaCl_2$, 25 mM Na HEPES, and 30 mM glucose at pH 7.4. All calcium imaging experiments were done at room temperature (~23°C). Baseline was acquired for 1 minute followed by stimulation (15 seconds) with an excitatory solution to activate low-voltage-activated $Ca^{2+}$ channels (i.e., R-type [Cav2.3] and T-type $Ca^{2+}$ channels [Cav3.1 to Cav3.3]) [containing 81.5 mM NaCl, 40 mM KCl, 2 mM $CaCl_2$, 2 mM $MgCl_2$, 25 mM HEPES, and 30 mM glucose at pH 7.4 and ~310 mOsm], or an excitatory solution to activate high-voltage-activated $Ca^{2+}$ channels (i.e., L-type [Cav1.1 to Cav1.4], N-type [Cav2.2], P/Q-type [Cav2.1], and R-type [CaV2.3]) [containing 32 mM NaCl, 90 mM KCl, 2 mM $MgCl_2$, 2 mM $CaCl_2$, 25 mM HEPES, and 30 mM glucose at pH 7.4 and ~310 mOsm]. Fluorescence imaging was performed with an inverted microscope, Nikon Eclipse T*i*-U (Melville, NY, USA), using objective Nikon Super Fluor ×10 0.50 and a photometrics cooled CCD camera CoolSNAP $ES^2$ (Roper Scientific, Tucson, AZ, USA) controlled by NIS Elements software (version 4.20; Nikon Instruments). The excitation light was delivered by a Lambda-LS system (Sutter Instruments, Novato, CA, USA). The excitation filters (340 ± 5 nm and 380 ± 7 nm) were controlled by a Lambda 10 to 2 optical filter change (Sutter Instruments). Fluorescence was recorded through a 505 nm dichroic mirror at 535 ± 25 nm. For minimization of photobleaching and phototoxicity, the images were taken every ~10 s during the time course of the experiment using the minimal exposure time that provided acceptable image quality. The changes in $[Ca^{2+}]_c$ were monitored by following a ratio of $F_{340}/F_{380}$, calculated after subtracting the background from both channels. All compounds were applied at 20 µM overnight for the initial screening.

**Constellation pharmacology.** These experiments were performed as described previously [18; 27; 28; 32], but with the following modifications. DRG neurons were loaded at 37°C with 3 µM Fura-2AM for 30 minutes in Tyrode's solution. After baseline acquisition (during perfusion with standard Tyrode's solution) for 1 minute, DRG neurons were sequentially stimulated for 15 seconds by the following receptor agonists (then washed with standard Tyrode's solution for 5 minutes between stimulation): 400 nM menthol, 50 µM histamine, 10 µM adenosine triphosphate (ATP), 200 µM allyl isothiocyanate (AITC), 1 mM acetylcholine (Ach), 100 nM capsaicin diluted in standard Tyrode's solution. At the end of the constellation pharmacology protocol, cell viability was assessed by depolarization-induced $Ca^{2+}$ influx using the excitatory solution for activating high-voltage-activated $Ca^{2+}$ channels. This process was automated using the ValveBank Controller (Automate Scientific; San Francisco, CA, USA). Fluorescence imaging was performed under the conditions described above for calcium imaging. A cell was classified as a "responder" if its maximum fluorescence ratio for 340 nm/380 nm exceeded 10% of its baseline value (calculated as the average fluorescence ratio during the 30 seconds immediately preceding application of the receptor agonist solution).

**Immunocytochemsitry.** Immunocytochemistry was performed on DRG neurons incubated with ***45*** (20µM) or its vehicle (DMSO) for 1 hour or 16 hours before cell fixation. A subset of cells was also treated with a proteasome inhibitor, lactacystin [12; 22]. Lactacystin treatment (10µM) was administered 16 hours before fixation, either concomitant to the 16-hours treatment with ***45*** or 15 hours before the 1-hour treatment with ***45***. Cells were fixed in 4% paraformaldehyde in PBS 20 minutes. The whole immunostaining process was conducted at room temperature. Permeabilization and saturation of non-specific binding sites was achieved by incubation in 3% BSA, 0.1% Triton X-100 in PBS for 1 hour. Cells were stained with an antibody against CaV2.2 (Origene, TA308673) diluted at 1/50 in PBS, 3% BSA for 3 hours then rinsed in PBS and incubated with the secondary antibody (Goat anti-Rabbit AlexaFluor 488, ThermoFisher, A11008) diluted at 1/250 in PBS, 3% BSA for 2 hours. Cells were thoroughly rinsed, and nuclei were counterstained with DAPI. Coverslips were mounted and stored at +4°C until image acquisition. Immunofluorescence micrographs were acquired using a Plan-Apochromat 20x (NA 0.8) objective on a Zeiss LSM880 confocal microscope operated by the Zen Black software (Zeiss). For quantitative analysis purposes, relevant parameters, such as laser intensities, pinhole size and detector gain, were kept constant between all experimental conditions. Analysis was performed on raw unmodified images. Images were analyzed using the free Icy software http://icy.bioimageanalysis.org)[6]. DRGs neurons were selected by thresholding on the CaV2.2, with manual adjustments when necessary. Mean immunofluorescence intensity for each whole cell was extracted (plug-in “Thresholder”). Membrane immunofluorescence was calculated by measuring the signal intensity in the area contiguous to the boundary of the cell. An annular region of interest (ROI) was obtained by rescaling the perimeter of each cell (plug-in “Rescale ROI”, rescaling factor: 0.8). The mean fluorescence intensity of the annular region between the perimeter and the inner curve was extracted and divided by the fluorescence intensity of the inner region (excluding the nuclear region, i. e. cytoplasmic region) to obtain a fluorescence ratio for each cell. Acquisition and analysis were performed by experimenters blinded to genotypes. For presentation purposes only, contrast was enhanced on immunofluorescence images. For statistical analysis, Kruskal-Wallis tests, followed by Dunn’s multiple comparisons tests were performed using the GraphPad Prism v7.04 software.

**Competition Radioligand Binding.** Competition radioligand binding experiments were performed as previously reported [26]. Membrane preparations of MOR-, DOR-, or KOR-CHO cells were combined with a fixed concentration of $^{3}$H-diprenorphine (MOR = 5.33 nM; DOR = 1.43 nM; KOR = 1.95 nM) and a concentration curve of competitor ligand. These reactions were miniaturized to a 200 μL volume in 96 well plates. The reaction proceeded at room temperature for 60 minutes. The reactions were terminated by rapid filtration through 96 well format GF/B filter plates (PerkinElmer) with cold water, washed, dried, and Microscint PS (PerkinElmer) added. The plates were read in a MicroBeta2 96 well format 6 detector scintillation counter (PerkinElmer). The data was normalized to the specific binding caused by $^{3}$H-diprenorphine alone (100%) or non-specific binding determined by a 10 μM concentration of known competitor ligand (0%; MOR = naloxone; DOR = SNC80; KOR = naloxone). $K_I$ values were calculated using the $IC_{50}$ of each competitor ligand and the previously established $K_D$ of $^{3}$H-diprenorphine in each cell line (GraphPad Prism 7.0).

**Whole-cell voltage-clamp electrophysiology.** Recordings were obtained from transfected HEK293 cells and acutely dissociated DRG neurons and performed at room temperature by using an EPC 10 Amplifier-HEKA. Electrodes were pulled from thin-walled borosilicate glass capillaries (Warner Instruments, Hamden, CT) with a P-97 electrode puller (Sutter Instrument, Novato, CA) such that final electrode resistances were 1–3 MΩ when filled with internal solutions. For HEK293 cells, external solution (in mM): 140 $Et_4N$ $MeSO_3$, 10 HEPES, and 5 $BaCl_2$ (pH 7.3); internal solution (in mM): 135 $CsMeSO_3$, 5 CsCl, 5 EGTA, 1 $MgCl_2$, 4 MgATP, and 10 HEPES (pH 7.3). For DRGs, external solution (in mM): (at ~315 mOsm, in mM): 110 N-methyl-D-glucamine (NMDG), 10 $BaCl_2$, 30 TEA-Cl, 10 HEPES, 10 glucose (pH 7.2 with KOH); internal solution: (at ~305 mOsm, in mM): 150 $CsCl_2$, 10 HEPES, 5 Mg-ATP, 5 BAPTA (pH 7.2 with KOH). To isolate current contributions of specific calcium channel subtypes, cells were treated with inhibitors of all other $Ca^{2+}$ channel subtypes; the following compounds were used: SNX482 (200 nM, R-type voltage-gated $Ca^{2+}$ channel blocker) [23], ω-conotoxin GVIA (500 nM, N-type voltage-gated $Ca^{2+}$ channel blocker) [8], ω-agatoxin (200 nM, P/Q-type voltage-gated $Ca^{2+}$ channel blocker) [16], TTA-P2 (1μM, T-type voltage-gated $Ca^{2+}$ channel blocker) [3], and Nifedipine (10 μM, L-type voltage-gated $Ca^{2+}$ channel blocker). Cells were subjected to current-voltage, inactivation, and use dependence protocols.

**Voltage-clamp protocols.** Analysis of $Ca^{2+}$ channel activation, as a function of current vs. voltage, as well as peak current density, which was typically observed near ∼0–10 mV and normalized to each cell's capacitance, was performed. In the current-voltage protocol, DRG neurons were held at resting membrane potential (RMP) for 5-ms before depolarization by 200-ms voltage steps from -70 mV to +60 mV in 10 mV increments. Currents were normalized to each cell's capacitance (pF). This allowed for collection of current density data to analyze activation of $Ca^{2+}$ channels as a function of current vs. voltage as well as peak current density. In the inactivation protocol, DRG neurons were held at -90 mV for 20 ms before depolarization by 1.5-s voltage steps from -100 mV to +10 mV in 10 mV increments, followed by a 20-ms pulse at 10 mV before returning to -90 mV for 20 ms. In the use dependence protocol, DRG neurons were held at -70 mV for 5-ms before depolarization by a 0 mV voltage step lasting 200-ms, followed by a return to -70 mV for 5-ms; this was repeated 30 times.

**Preparation of spinal cord slices.** As described previously [34], young rats (postnatal 12-21 days) were deeply anesthetized with isoflurane. For spinal nerve blocking, 0.3 mL of 2% lidocaine was injected to both sides of L4 to 5 lumbar vertebrae. Laminectomy was performed from mid-thoracic to low lumbar levels, and the spinal cord was quickly moved to cold modified artificial cerebrospinal fluid (ACSF) oxygenated with 95% $O_2$ and 5% $CO_2$. The ACSF contained (in mM): 80 NaCl, 2.5 KCl, 1.25 $NaH_2PO_4$, 0.5 $CaCl_2$, 3.5 $MgCl_2$, 25 $NaHCO_3$, 75 sucrose, 1.3 ascorbate, 3.0 sodium pyruvate, at pH 7.4 and ~310 mOsm. Transverse 350-mm thick slices were obtained by a vibratome (VT1200S; Leica, Nussloch, Germany). Slices were then incubated for at least 1 hour at RT in an oxygenated recording solution containing (in mM): 125 NaCl, 2.5 KCl, 2 $CaCl_2$, 1 $MgCl_2$, 1.25 $NaH_2PO_4$, 26 $NaHCO_3$, 25 D-glucose, 1.3 ascorbate, 3.0 sodium pyruvate, at pH 7.4 and ~320 mOsm. The slices were then positioned in a recording chamber and continuously perfused with oxygenated recording solution at a rate of 3 to 4 mL/min before electrophysiological recordings at RT.

**Electrophysiological recording in spinal cord slices by whole-cell patch clamp.** Substantia gelatinosa neurons were visualized and identified in the slices by means of infrared differential interference contrast video microscopy on an upright microscope (FN1; Nikon, Tokyo, Japan) equipped with a 3 40/0.80 water-immersion objective and a charge-coupled device camera. Electrodes were pulled from thin-walled borosilicate glass capillaries (Warner Instruments, Hamden, CT) with a P-97 electrode puller (Sutter Instrument, Novato, CA) such that final electrode resistances were 6–10 MΩ when filled with internal solutions. The internal solution contained the following (in mM): 120 potassium gluconate, 20 KCl, 2 MgCl 2, 2 Na 2 -ATP, 0.5 Na-GTP, 20 HEPES, 0.5 EGTA, with pH 7.28 and ~310 mOsm. Membrane potential was held at -60 mV using PatchMaster software and a dual channel EPC10-HEKA amplifier (Lambrecht, Germany).

Whole-cell configuration was obtained in voltage-clamp mode. To record spontaneous excitatory postsynaptic currents (sEPSCs), bicuculline methiodide (10 μM) and strychnine (1 μM) were added to the recording solution to block γ-aminobutyric acid-activated and glycine-activated currents. Hyperpolarizing step pulses (5 mV in intensity, 50 milliseconds in duration) were periodically delivered to monitor the series resistance (15-25 MΩ), and recordings were discontinued if the series resistance changed by more than 20%. For each neuron, sEPSCs were recorded for a total duration of 2 minutes. Currents were filtered at 3 kHz and digitized at 5 kHz. Data were further analyzed by the Mini-Analysis Program (Synatosoft Inc, NJ, USA) to provide spreadsheets for the generation of cumulative probability plots. The amplitude and frequency of sEPSCs were compared between neurons from animals in control and compound **45** groups.

**Calcitonin gene–related peptide release from lumbar slices.**

Rats were deeply anesthetized with 5% isofluorane and then decapitated. Two vertebral incisions (cervical and lumbar) were made in order to expose the spinal cord. Pressure was applied to a saline-filled syringe inserted into the lumbar vertebral foramen, and the spinal cord was extracted. Only the lumbar region of the spinal cord was used for the CGRP release assay. Baseline treatments (#1 and #2) involved bathing the spinal cord in Tyrode's solution. The excitatory solution consisting of 90 mM KCl (also used for the calcium imaging experiments) was paired with the treatment for fraction #4. These fractions (10 minutes, 400 μL each) were collected for measurement of CGRP release. Samples were immediately flash frozen and stored in a -20℃ freezer. Compound *45* (20 μM) or vehicle (0.9% saline) was added to the pretreatment and co-treatment fractions (#3 and 4). The concentration of CGRP released into the buffer was measured by enzyme-linked immunosorbant assay (Cat# 589001, Cayman Chemical, Ann Arbor, MI).

**Synapse enrichment and fractionation.** Adult rats were killed by isofluorane overdose and decapitation, the spinal cords dissected and the lumbar dorsal horn collected. Only the dorsal horn of the spinal cord was used as this structure contains the synapses arising from the DRG. Synaptosomes were isolated as described previously [24]. Fresh tissues were homogenized in ice-cold Sucrose 0.32M, HEPES 10 mM, pH 7.4 buffer. The homogenates were centrifuged at 1000xg for 10 min at 4°C to pellet the insoluble material. The supernatant was harvested and centrifuged at 12000xg for 20 min at 4°C to pellet a crude membrane fraction. The pellet was then re-suspended in a hypotonic buffer (4 mM HEPES, 1 mM EDTA, pH 7.4) and the resulting synaptosomes pelleted by centrifugation at 12000xg for 20 min at 4°C. The synaptosomes were then incubated in 20 mM HEPES, 100 mM NaCl, 0.5% triton X, pH= 7.2) for 15 min on ice and centrifuged at 12000xg for 20 min at 4°C. The supernatant was considered as the non-postsynaptic density (non-PSD) membrane fraction, sometimes referred to as the triton soluble fraction. The pellet containing the postsynaptic density fraction (PSD) was then solubilized in RIPA buffer (50mM Tris-HCl, pH 7.4, 50 mM NaCl, 2 mM $MgCl_2$, 1% [vol/vol] NP40, 0.5% [mass/vol] sodium deoxycholate, 0.1% [mass/vol] SDS) as described previously [9]. All buffers contained protease inhibitors (Cat# B14002; Bimake, Houston, TX) and phosphatase inhibitors (Cat# B15002, Bimake). The integrity of non-PSD and PSD fractions was verified by immunoblotting for PSD95, which was enriched in PSD fraction, and synaptophysin which was enriched in non-PSD fraction. All buffers were supplemented with protease and phosphatase inhibitor cocktails. Protein concentrations were determined using the BCA protein assay.

**Immunoblot preparation and analysis**. Indicated samples were loaded on 4-20% Novex® gels (Cat# EC60285BOX, Thermo Fisher Scientific, Waltham, MA). Proteins were transferred for 1h at 120 V using TGS buffer (25mM Tris pH=8.5, 192mM glycine, 0.1% (mass/vol) SDS), 20% (vol/vol) methanol as transfer buffer to polyvinylidene difluoride (PVDF) membranes 0.45µm (Cat# IPVH00010, Millipore, Billerica, MA), pre-activated in pure methanol. After transfer, the membranes were blocked at room temperature for 1 hour with TBST (50 mM Tris-HCl, pH 7.4, 150 mM NaCl, 0.1 % Tween 20), 5% (mass/vol) non-fat dry milk, then incubated separately with the primary antibodies CaV2.2 (Cat# TA308673, Origene, Rockville, MD), Synaptophysin (Cat# MAB5258, Thermofisher scientific, San Diego, CA), PSD95 (Cat# MA1-045, Thermofisher scientific) or Flotilin (Cat# F1180, Sigma, St. Louis, MO) in TBST, 5% (mass/vol) BSA, overnight at 4°C. Following incubation in horseradish peroxidase-conjugated secondary antibodies from Jackson immunoresearch, blots were revealed by enhanced luminescence (WBKLS0500, Millipore, Billerica, MA) before exposure to photographic film. Films were scanned, digitized, and quantified using Un-Scan-It gel version 6.1 scanning software by Silk Scientific Inc.

**Indwelling intrathecal (i.t.) catheter.** Rats were anesthetized (ketamine or xylazine anesthesia, 80/12 mg/kg intraperitoneally (i.p.) injected; Sigma) and placed in a stereotaxic head holder. The cisterna magna was exposed and incised. An 8-cm catheter (PE-10; Stoelting, Wood Dale, IL) was implanted as previously reported, terminating in the lumbar region of the spinal cord. [33] Catheters were sutured (using 3-0 silk sutures) into the deep muscle and externalized at the back of the neck; skin was closed with autoclips, and other surgeries were performed after a 5 to 7-day recovery period.

**Measurement of thermal withdrawal latency**. The method of Hargreaves et al. was used [10]. Rats (naïve or following experimentally-induced pain (see below)) were acclimated within Plexiglas enclosures on a clear glass plate maintained at 30°C. A radiant heat source (high-intensity projector lamp) was focused onto the plantar surface of the hind paw. When the paw was withdrawn, a motion detector halted the stimulus and a timer. A maximal cut-off of 33.5 s was used to prevent tissue damage.

**Measurement of allodynia.** To assess tactile allodynia (i.e., a decreased threshold to paw withdrawal after probing with normally innocuous mechanical stimuli), rats' paw withdrawal threshold in response to probing with a series of fine (von Frey) calibrated filaments was measured. Rats were held in suspended wire mesh cages, and each von Frey filament was perpendicularly applied to the plantar surface of the rats' paw. Withdrawal threshold was determined by sequentially increasing and decreasing the stimulus strength (the 'up-and-down' method), and data were analyzed with Dixon's nonparametric method, as described by Chaplan et al. [2; 7], and expressed as the mean withdrawal threshold.

**Paw incision model of postoperative pain.** A rodent model of surgical pain was generated by plantar incision as previously described [1]. Male Sprague–Dawley rats were anesthetized with isoflurane vaporized through a nose cone. The plantar aspect of the left hind paw was scrubbed with betadine and 70% alcohol three times. A 1-cm long incision, starting 0.5 cm from the heel and extending toward the toes, was made with a number 11 blade, through the skin and fascia of the plantar aspect of the left hind paw including the underlying muscle. The plantaris muscle was then elevated and longitudinally incised, leaving the muscle origin and insertion intact. After hemostasis with gentle pressure, the skin was closed with 2 mattress sutures of 5-0 nylon on a curved needle. Rats received an injection of gentamicin (1mL/kg of 8 mg/mL solution, injected subcutaneously) and were allowed to recover from anesthesia before returning to their home cage. Sham rodents were anesthetized, and the left hind paw was scrubbed with betadine and 70% ethanol three times, but no incision was made. Following surgery, rodents were allowed to recover for 24 hours; then, both paw withdrawal thresholds and paw withdrawal latencies were measured.

**Spinal nerve ligation model of neuropathic pain.** Nerve ligation injury produces signs of neuropathic dysesthesias, including tactile allodynia and thermal hypersensitivity. All nerve operations occurred 5 days after intrathecal catheter implantation. Rats1 were anesthetized with 2% isoflurane in $O_2$ anesthesia delivered at 2 L/min. The skin over the caudal lumbar region was incised, and the muscles retracted. The L5 and L6 spinal nerves were exposed, carefully isolated, and tightly ligated with 4-0 silk distal to the DRG, without limiting the use of the left hind paw of the animal. All animals were allowed

7 days to recover before any behavioral testing. Any animal exhibiting signs of motor deficiency was killed.

**Elevated plus maze test of anxiety-related behaviors.** One hour after intraperitoneal (i.p.) injection of ***45*** or its vehicle, mice (CD1) were placed at the center of the elevated plus-maze, where the two arms intersect and allowed to explore the maze freely for 10 minutes. The apparatus dimensions were as follow: arm length: 35 cm; arm width: 5 cm; wall height for closed walls: 15 cm. The whole apparatus is elevated 50 cm above ground. Animals movements were recorded by a camera (C310, Logitech) placed above the apparatus and tracked and analyzed using the Any-MAZE software (v6.05, Stoelting). Entry into a zone was defined as 70% of the animal's body being in the zone. The main parameter analyzed was the time spent into the open arms of the apparatus.

**Acute pain models.** The tail flick response was evoked by a light beam (irradiated heat) or by immersing one third of the tail in water at 52 °C. In the hot plate test, mice were placed on a plate (52 °C), and responses were counted when the mouse first licked the hind paws and jumped from the plate (cut-off time, 10 s and 20 s, respectively).

**HIV-induced sensory neuropathy.** Mechanical allodynia is produced by i.t. administration of the human immunodeficiency virus-1 (HIV-1) envelope glycoprotein, gp120 [15]. Seven days after implantation of an i.t. catheter, baseline behavioral measurements were obtained and then rats were randomly assigned to two groups. On days 10, 12 and 14, rats were injected i.t. with 300 ng of gp120 (Cat#4961, HIV-1 BaL gp120 recombinant protein, NIH-AIDS Reagent program) in a final volume of 20 μL in 0.9% saline and 0.1% BSA.

**Chemotherapy-induced peripheral neuropathy via Paclitaxel.** Rats were given paclitaxel (Cat# P-925-1, Goldbio, Olivette, MO) based on the protocol described by Polomano et al. [25]. In brief, pharmaceutical-grade paclitaxel (Taxol) was re-suspended at a concentration of 2 mg/ml in 30% 1:1 Cremophor EL: ethanol, 70 % Saline and given to the rats at 2 mg/kg intraperitoneally (i.p.) every other day for a total of 4 injections (days 0, 2, 4, and 6), resulting in a final cumulative dose of 8 mg/kg. No abnormal spontaneous behavioral changes in the rats were noted during or after the treatment. Animals developed mechanical hyperalgesia within 10 days after the first paclitaxel injection.

**In vivo transfection of CRISPR plasmids for Cas9-mediated editing of *Nf1*.** As described before [17; 19; 21], NF1-related pain was induced by CRISPR/Cas9 mediated editing of the exon 39 of the gene *Nf1***.** The specific gRNA sequence was cloned in the plasmid pL-CRISPR-EFS-tRFP [11] and was validated in [17]. Lentivral particle were produced by Viracore (UCSF) and provided concentrated at a titer higher than $10^7$ infectious particles per ml. at day 0, 5 μl of the lentivral concentrate was injected intrathecally in male Sprague dawley rats. After 10 days, rats developed thermal hyperalgesia as described before [17; 19; 21].

**Rotarod**

Rats were trained to walk on a rotating rod (10 rev/min; Rotamex 4/8 device) with a maximal cutoff time of 180 seconds. Training was initiated by placing the rats on a rotating rod and allowing them to walk until they either fell off or 180 seconds was reached. This process was repeated 6 times. Prior to treatment (intracerebroventricular (i.c.v.) injection of compound *45* or saline), the rats were run once on a moving rod in order to establish a baseline value. Assessment consisted of placing the rats on the moving rod and timing until either they fell off or reached a maximum of 180 seconds.
